# Biologically-derived neoproteoglycans for profiling protein-glycosaminoglycan interactions

**DOI:** 10.1101/2022.03.04.482917

**Authors:** Ryan N. Porell, Julianna L. Follmar, Sean C. Purcell, Bryce Timm, Logan K. Laubach, David Kozirovskiy, Bryan E. Thacker, Charles A. Glass, Philip L. S. M. Gordts, Kamil Godula

## Abstract

Glycosaminoglycans (GAGs) are a class of highly negatively charged membrane associated and extracellular matrix polysaccharides involved in the regulation of myriad biological functions, including cell adhesion, migration, signaling and differentiation, among others. GAGs are typically attached to core proteins, termed proteoglycans (PGs), and can engage >500 binding proteins, making them prominent relays for sensing external stimuli and transducing cellular responses. However, their unique substructural protein-recognition domains that confer their binding specificity remain elusive. While the emergence of glycan arrays has rapidly enabled the profiling of ligand specificities of a range of glycan-binding proteins, their adaptation for the analysis of GAG-binding proteins has been considerably more challenging. Current GAG-microarrays primarily employ synthetically defined oligosaccharides, which capture only a fraction of the structural diversity of native GAG polysaccharides. Augmenting existing array platforms to include GAG structures purified from tissues or produced in cells with engineered glycan biosynthetic pathways may significantly advance the understanding of structure-activity relationships in GAG-protein interactions. Here, we demonstrate an efficient and tunable strategy to mimic cellular proteoglycan architectures by conjugating biologically-derived GAG chains to a protein scaffold, defined as neoproteoglycans (neoPGs). The use of a reactive fluorogenic linker enabled real-time monitoring of the conjugation reaction efficiency and tuning of the neoPG valency. Immobilization of the reagents on a 96-well array platform allowed for efficient probing of ligand binding and enzyme substrate specificity, including growth factors and the human sulfatase 1. The neoPGs can also be used directly as soluble probes to evaluate GAG-dependent growth factor signaling in cells.

**Graphical Abstract:** 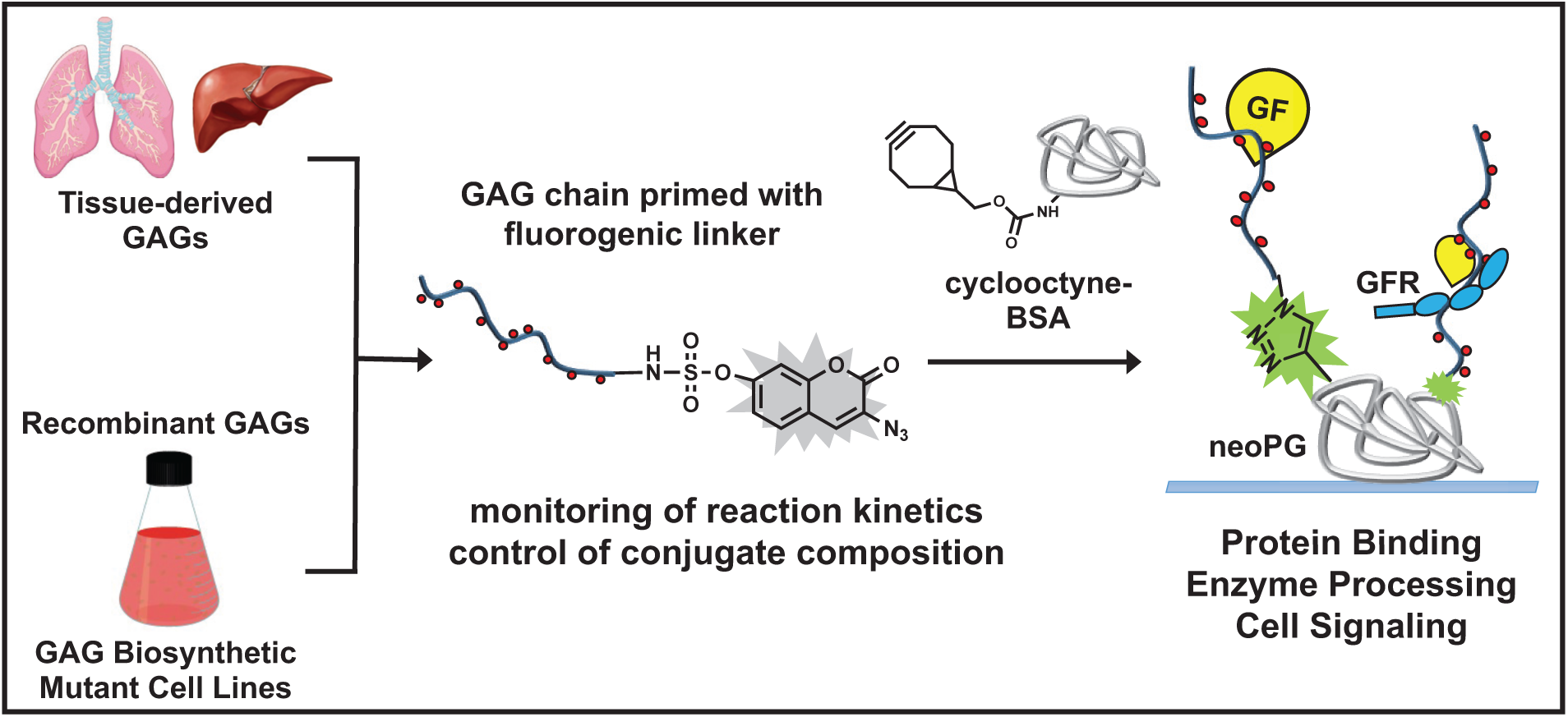

## Introduction

Proteoglycans (PGs) are abundant on cell surfaces and in the extracellular matrix, where they serve a myriad of biological functions spanning from regulation of growth factor binding and signaling^1,2,3^ to tissue development and organ function.^4,5^ They also contribute to pathophysiological processes, including aging and associated disease,^6^ immunological responses,^7,8^ and they serve as receptors for infectious agents, such as the Herpes simplex viruses^9^ and the SARS-CoV-2 virus,^10^ among others. Key structural components of PGs, which define their interactions with other proteins, are sulfated glycosaminoglycan (GAG) polysaccharide chains appended to the protein core via a glycosidic bond to serine and threonine residues. Sulfated GAGs can be classified according to their monosaccharide composition as heparan sulfate (HS), chondroitin sulfate (CS), dermatan sulfate (DS), and keratan sulfate (KS). Hyaluronan (HA), which is the last member of the GAG family, lacks sulfation and is not attached to proteins (Figure 1).

**Figure 1:**
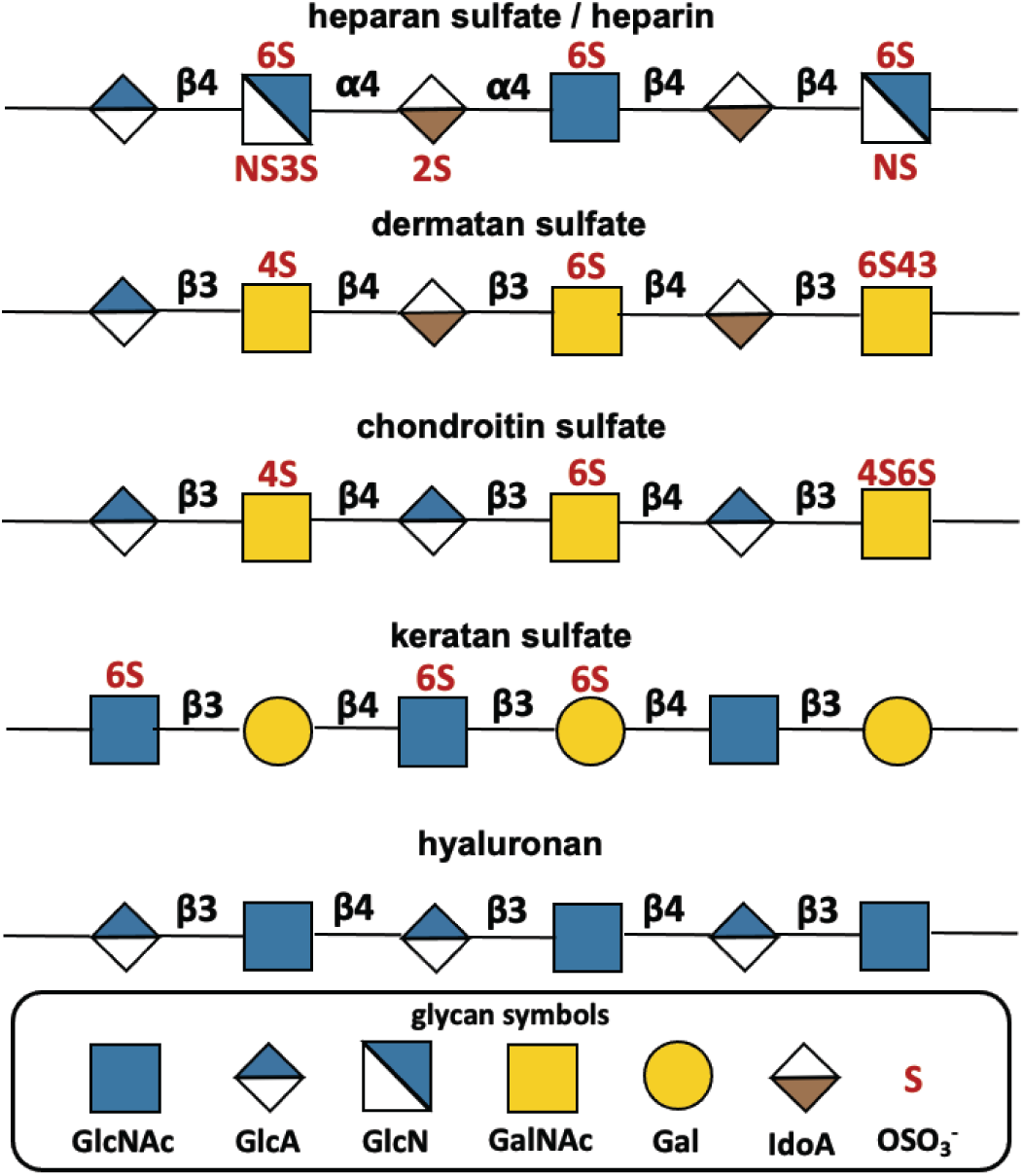
Common structures of GAGs using the glycan symbol nomenclature. Possible sulfation sites are displayed in red text above or below their respective monosaccharide.

The biological specificity of GAGs is established during their biosynthesis through a non-templated process via a sequence of enzymatic modifications, which elongate the individual polysaccharide chains and install negatively charged sulfate groups. This results in structurally complex sulfation patterns organized in domains along the polysaccharides that provide high-affinity binding sites for proteins.^11,12^ The structure-function relationships for GAGs remain poorly defined, due to challenges in isolation and structural characterization of GAGs from biological samples, as well as difficulties with producing structurally defined GAG polysaccharides synthetically.

Since their inception in 2002,^13,14,15^ glycan arrays have become broadly adapted as a high throughput glycomics tool to profile ligand specificity of glycan binding proteins.^16^ The glycan array developed by the National Center for Functional Glycomics^17^ now contains more than 1,000 unique structures representing *N-* and *O*-linked glycans and is considered the gold standard for the field. Parallel pioneering efforts to establish GAG arrays using synthetic HS^18^ and CS^19^ oligosaccharides provided early insights into GAG structure-function relationships. Advances in chemoenzymatic synthesis of GAG oligosaccharides have significantly accelerated these efforts in recent years.^20,21,22^ Upwards of 95 synthetic HS structures are now available in an array format,^23^ which is still a significant shortfall from the total amount of GAG structures that are found naturally.

Complementing bottom-up synthetic efforts are arrays consisting of biologically-derived GAGs representing the structural complexity of the natural polysaccharides, however, these efforts faced major roadblocks due to difficulties in purification and separation of GAGs into structurally distinct populations.^24^ Recent advances in generating glycosylation mutant cell lines through systematic genetic manipulation of GAG biosynthesis are alleviating this limitation by providing access to increased quantities of compositionally-defined bioengineered GAGs.^25,26^

A technical challenge in generating native and bioengineered GAG arrays is the optimization of GAG immobilization onto surfaces. Commonly employed approaches include electrostatic adsorption of the polyanionic glycans onto positively charged poly(lysine)-coated surfaces^27^ or precipitation on plastic supports using ammonium chloride.^28^ These methods unequally sequester biologically active sulfated domains and influence protein binding. Alternatively, covalent conjugation of GAGs via their reducing ends to amine-, aminooxy-, or hydrazine-functionalized surfaces allows for glycan extension away from the surface, enhancing chain presentation.^29,30^ However, chain grafting efficiency is generally low and varies based on the length and charge of the GAG structure. The inability to characterize the GAGs presentations after immobilization introduces a degree of uncertainty making it difficult to perform comparative analysis of protein binding specificity.

Here, we introduce semi-synthetic neoproteoglycan (neoPG) reagents with a defined molecular architecture to permit comparative analysis of GAG-protein binding interactions. The neoPG has GAG polysaccharides chains end-conjugated to a carrier bovine serum albumin (BSA) protein, improving on existing reductive amination protocols.^31^ We employ the strain-promoted alkyne-azide cycloaddition (SPAAC) reaction^32^ with a reactive fluorogenic linker to generate the glycoconjugates. The strategy enables efficient coupling with real-time monitoring of GAG conjugation and quantification of the neoPG composition. This novel method is suitable for all members of the GAG family, including tissue-derived and bioengineered polysaccharides and the reagents can be immobilized in an ELISA format to analyze GAG-binding protein interactions or used as soluble reagents to evaluate signaling activity in cells.

## Results and Discussion

### Generation of neoProteoglycans (neoPGs)

To generate versatile reagents suitable for analysis of GAG-interactions in analytical assays as well as biological assays, we developed a chemical approach for merging polysaccharides with a protein carrier without disrupting ligand-binding domains. Covalent GAG attachment to the protein backbone mimics the organization of native PGs and provides control over GAG valency and presentation both in solution and after immobilization on surfaces. However, macromolecular assemblies of the highly sulfated GAG polysaccharides with other macromolecules, including proteins, necessitates efficient and high-yielding bioconjugation chemistries. This is particularly challenging for conjugation of GAGs from biological samples, which can only be isolated in limited amounts or otherwise become costly. To address these challenges, we have developed a fluorogenic bioorthogonal linker strategy for attaching GAG chains through their reducing ends to a BSA protein carrier. In this process, the GAG chains are furnished with a novel azido-coumarin linker that produces fluorescent light emission (λ _ex/em_ = 393/477 nm) upon further conjugation with alkynes. This strategy enables direct monitoring and quantification of the coupling reaction (Figure 2b). The linker is introduced via the sulfur (IV) fluoride exchange (SufEx)^33^ reaction between 3-azidocoumarin-7-sulfonyl fluoride (**ACS-F**) and amine-terminated GAG chains. The amine groups either originate from amino acid residues retained after GAG release from proteoglycans by pronase digestion or are introduced quantitatively to the hemiacetal end of β-eliminated chains by treatment with *N*-methylaminooxy propylamine (Figure S6).^34,35^ After removal of excess **ACS-F** linker by size exclusion chromatography and dialysis, the chemically primed GAGs (**ACS-GAGs**) were conjugated to a bicyclo[6.1.0]nonyne-modified BSA protein (**BCN-BSA**) via the strain-promoted azide-alkyne cycloaddition (SPAAC)^36^ to generate neoPGs (Figure 2a). Fluorescence readout from the fluorogenic linker provided conjugation kinetics and stoichiometry of the resulting neoPGs, which was confirmed through a combination of BCA and carbazole assays to determine the respective BSA and GAG content after removal of unreacted GAG chains by dialysis. The fluorescence from triazole-coumarin sulfonyl-BSA (**TCS-BSA**) conjugate produced by reacting **BCN-BSA** with the **ACS-F** linker alone was used to calibrate the measurement and to establish the maximal number of conjugation sites (∼16 cyclooctynes per BSA, Figure S5) on the protein (Figure 2b). Using this approach, we prepared neoPG conjugates using polysaccharides representing the main classes of GAGs. These included commercially available heparin (Hep, a highly sulfated form of HS), HA, and bovine cartilage CS as well as HS, CS and KS isolated from pig lung and mouse liver tissues (Table S2). Under optimized conditions, **BCN-BSA** (∼1 nM) was reacted with **ACS-GAGs** (∼10 equiv. per BSA) in PBS buffer at ambient temperature for 20 hrs. Each neoPG was assigned a descriptor **GAG**_**x**_**-BSA**, where x designates the number of GAG chains per BSA molecule. The conjugation process was efficient, and the maximum number of GAG chains introduced into the neoPGs ranged from x ∼ 6-8 for HS, KS and HA and x ∼ 8-12 for CS. Both the size and charge of the polysaccharides likely contribute to the overall efficiency of the conjugation process; however, we did not observe any noticeable trends (Figure 2c and Table S2). The composition of the conjugates with respect to GAG chain valency can be tuned by controlling the reagent stoichiometry or reaction time.

**Figure 2:**
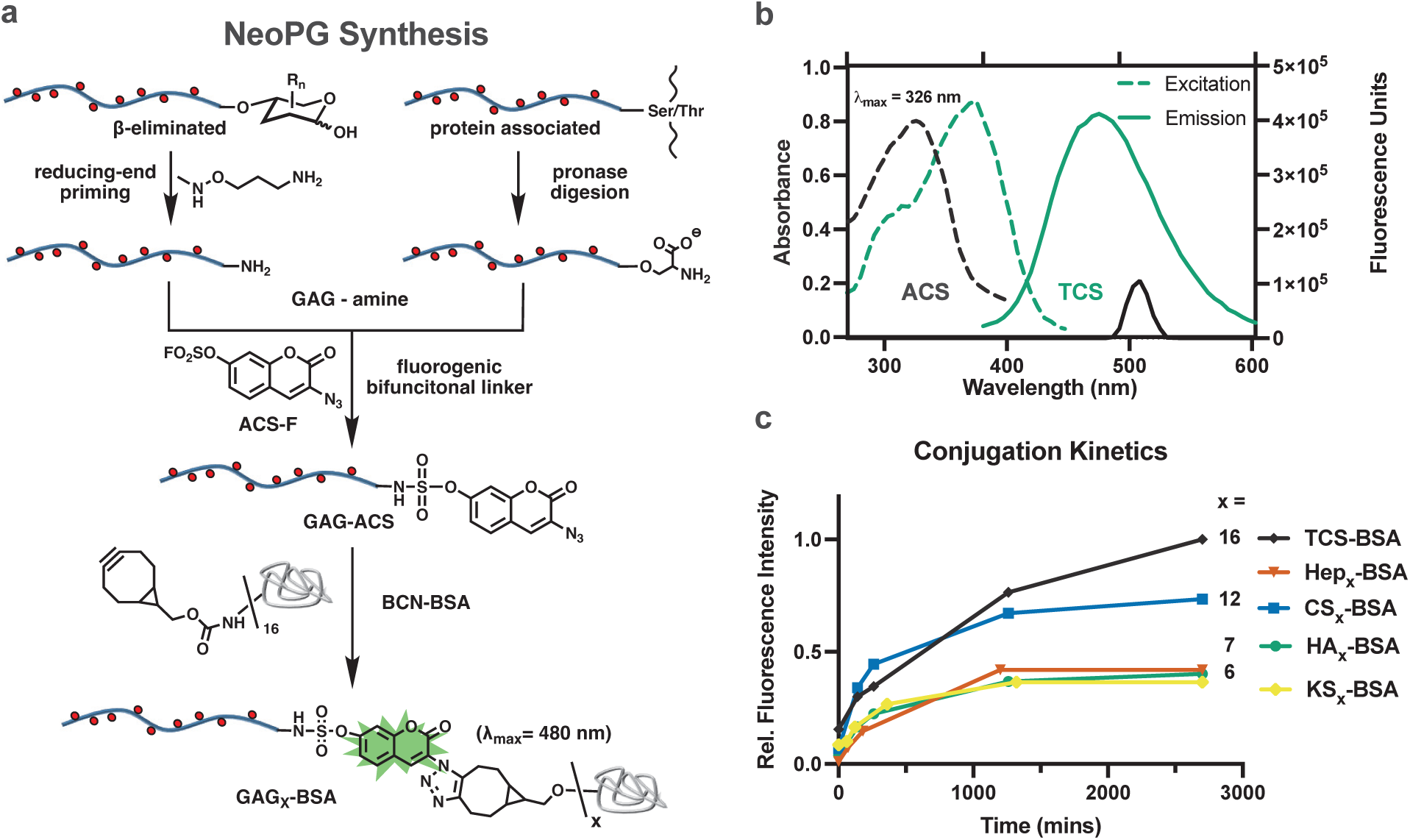
Preparation of HS, CS, KS, & HA neoPGs. **a)** Workflow for neoPG synthesis using recombinant or tissue purified GAGs. Fluorogenic bifunctional azidocoumarin sulfonyl fluoride (**ACS-F**) linker was conjugated via the sulfur (IV) fluoride exchange reaction to GAG chains primed at their reducing end with reactive amines. Subsequent strain-promoted azide-alkyne cycloaddition (SPAAC) reaction with cyclooctyne-functionalized BSA (**BCN-BSA**) furnished the desired neoPGs. Fluorescence signal produced upon conversion of the **ACS** handle into triozolylcoumarin sulfonamide (**TCS**) was used to monitor the progress of the conjugation reaction. **b)** Absorbance (dashed) and fluorescence (solid) spectra of quenched **ACS-F** (black) and unquenched **TCS-BSA** (green). **c)** Fluorescence emission (393 nm Ex./477 nm Em.) was used to monitor conjugation kinetics for Hep (red), CS (blue), HA (green), and KS (yellow) neoPGs. GAG chain valency of the resulting neoPGs was determined using **TCS-BSA** (black) as a standard.

### NeoPGs allow for controlled surface GAG displays

The neoPG reagents can be readily immobilized onto high-binding 96-well microtiter plates through adsorption via their BSA protein core (Figure 3). The density of the immobilized GAG chains defines the avidity of the protein interactions with the surface. Accordingly, this was reflected in the binding response for the fibroblast growth factor 1 (FGF1), a well characterized HS-binding growth factor, to plates treated with increasing concentrations of **Hep**_**7**_**-BSA** (20-500 ng/well, Figure 3a). Likewise, the valency of the neoPG conjugates can influence the avidity of protein binding. By controlling the **Hep-ACS** to **BCN-BSA** ratio during the conjugation reaction, we prepared neoPG conjugates displaying 1, 2 or 4 Hep chains. The valency-variant **Hep**_**x**_**-BSA** neoPGs were arrayed on plates at sub-saturation concentration (100 ng/well, Figure 3a) and probed for binding of FGF1. When normalized to BSA concentration, we observed increasing FGF1 binding reflective of the overall amount of heparin introduced to each well (Figure 3b, blue bars). When analyzed based on Hep concentration we observed a significant improvement in FGF1 binding between the monovalent and divalent neoPGs, **Hep**_**1**_**-BSA** and **Hep**_**2**_**-BSA**, with no further increase in avidity for the tetravalent neoPG, **Hep**_**4**_**-BSA** (Figure 3b, green bars).

**Figure 3:**
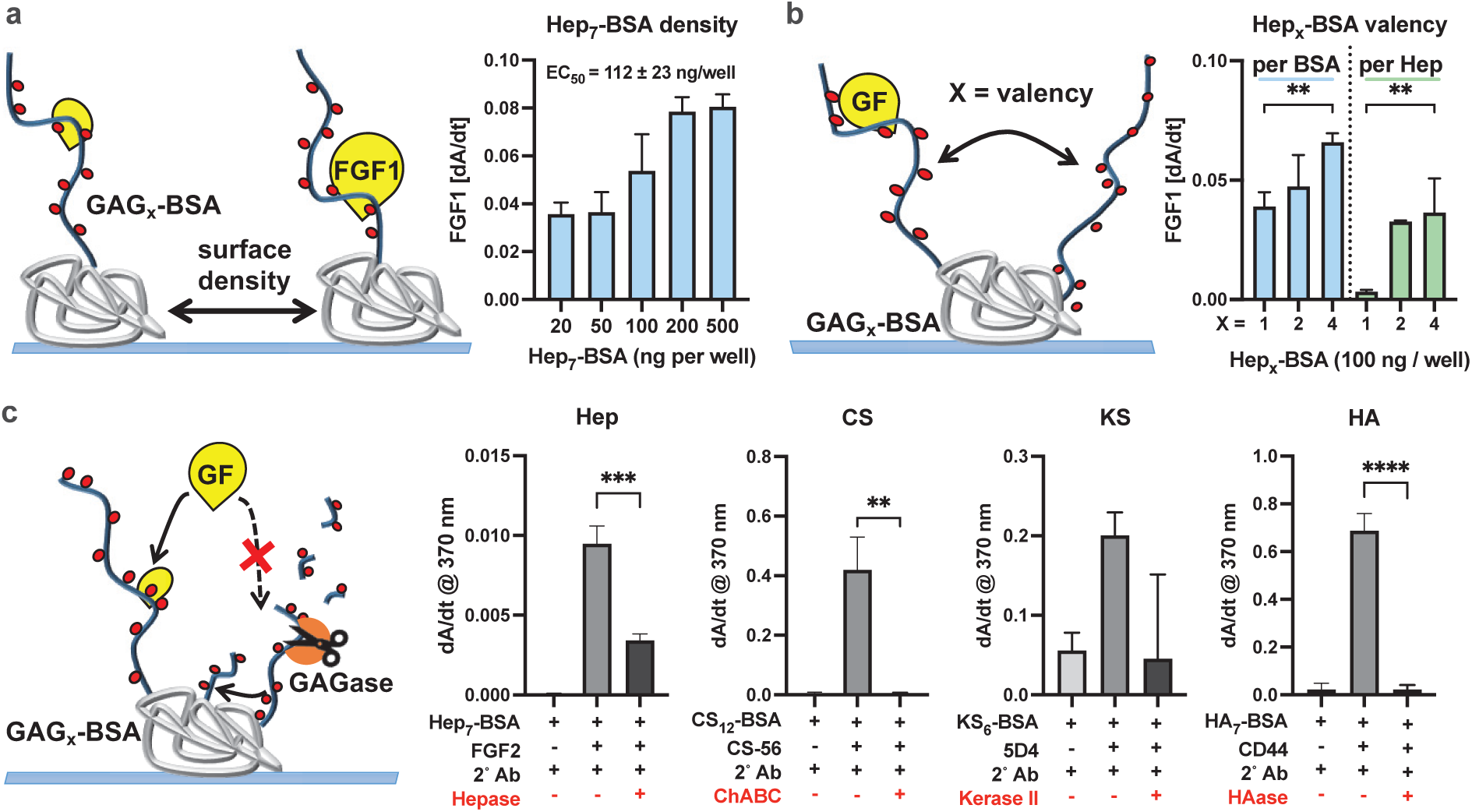
Construction and validation of neoPG arrays in ELISA format. **a)** FGF1 binding response to increasing surface density of **Hep**_**7**_**-BSA. b)** FGF1 binding response to increasing Hep-BSA valency normalized to BSA (blue bars) and heparin (green bars) concentration. **c)** Effects of neoPG degradation by specific GAG-lyases (red) on protein binding. **Hep**_**7**_**-BSA** treated with heparin lyases and probed for FGF2 binding. **CS**_**12**_**-BSA** treated with chondroitinase ABC and probed for CS-56 antibody binding. **KS**_**6**_**-BSA** treated with keratanase II and probed for 5D4 antibody binding. **HA**_**7**_**-BSA** treated with hyaluronidase and probed for CD44 binding. (Bar graphs represent n = 3 replicates, *p*-values were determined using student’s t-test, \*\**p* < 0.01, \*\*\**p*< 0.001, \*\*\*\**p*< 0.001,).

To confirm that the neoPG conjugates retained the protein binding specificities of their parent GAGs after immobilization, we arrayed **Hep**_**x**_**-, CS**_**x**_**-, KS**_**x**_**-**, and **HA**_**x**_**-BSA** (100 ng/well) and tested them with proteins and antibodies with known binding activities against these GAGs, i.e., FGF2, CS-56, 5D4 and CD44, respectively (Figure 3c). Degradation of the GAG chains with HS-, CS-, KS-and HA-specific glycosidase enzymes reduced the binding of these proteins to the neoPGs, further confirming glycan-dependent interactions. The immobilization of neoPGs through their core BSA proteins thus allows for controlled presentation of GAG chains with respect to valency and surface density to evaluate protein binding and in a format accessible to GAG-processing enzymes.

### Bioengineered recombinant HS neoPG arrays to analyze GF binding and sulfatase activity

Recent progress in systematic genetic manipulation of cellular HS biosynthesis has provided access to cell lines producing recombinant HS (rHS) structures with differences in overall level, type, and organization of sulfation (Figure 4a). These reagents can provide new insights into the structure-activity relationships in GAG-protein interactions and help define the substrate specificities of GAG-processing enzymes. Using our optimized conjugation method, we have converted a set of rHS polysaccharides produced in genetically engineered Chinese Hamster Ovary (CHO) cells and mastocytoma cells^37^ into neoPGs. All conjugates were prepared using the same **rHS-ACS** to **BCN-BSA** stoichiometry (20 : 1) and the reactions were perfomed for 48 hrs to maximize the extent of conjugation. The resulting neoPG conjguates were purified by dialysis to remove unreacted **rHS-ACS** and their valencies were determined by fluorescence reading (Figure S9). The panel of bioengineered neoPGs included rHS structures with overall sulfation levels similar to those found in native HS on cell surfaces and in tissues (i.e., rHS01, rHS02 and rHS08, Figure 4a). Compared to rHS01, the rHS02 lacked 2-*O*-sulfation, while rHS08 presented additional 3-*O*-sulfation. Also included were highly sulfated structures rHS09 and rHS29, which are similar to heparin but differed in their level of 2-O-sulfation and both lacked 3-*O*-sulfation (Figure 4a).

**Figure 4:**
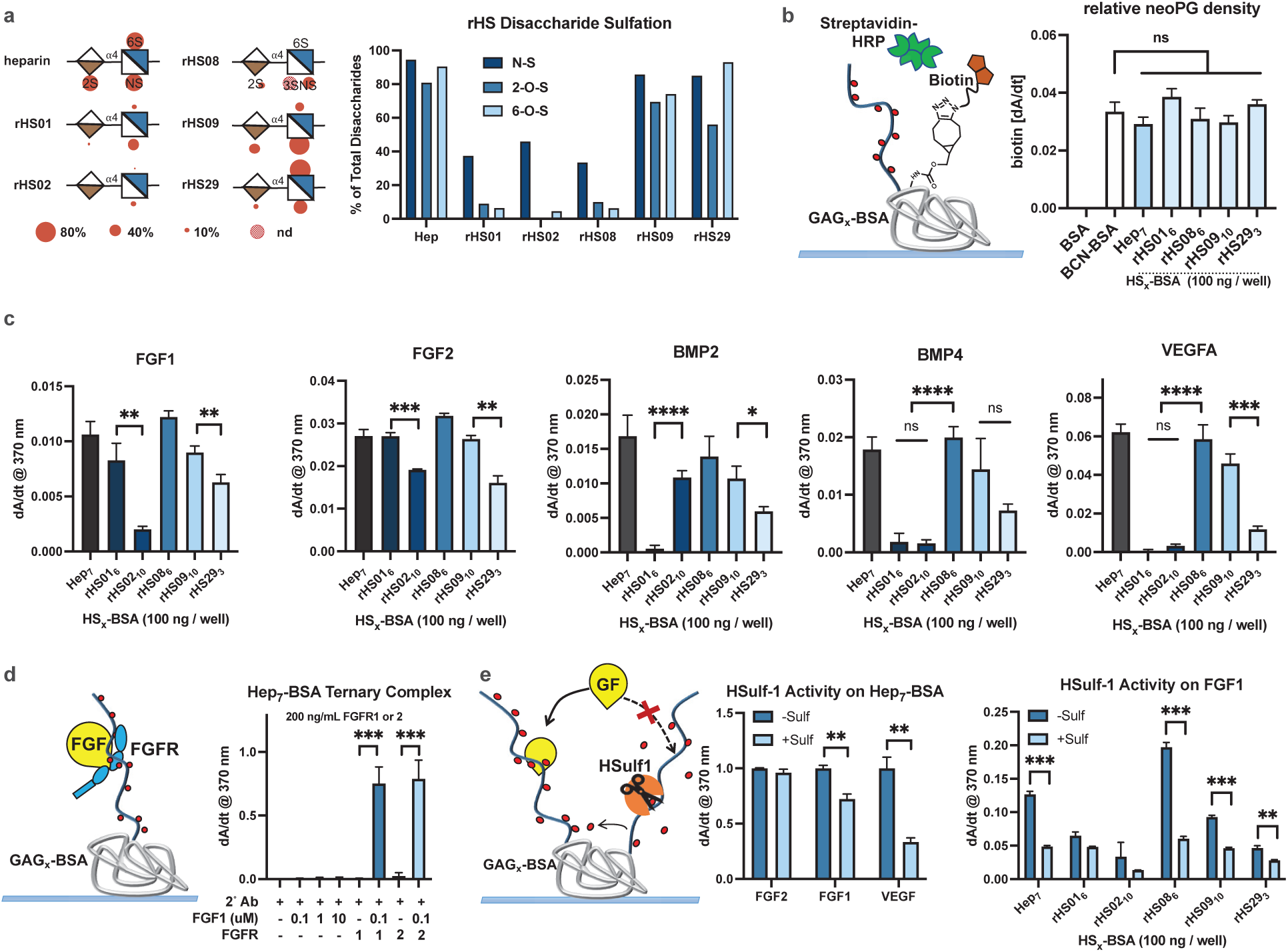
Analysis of protein interactions with rHS neoPGs via ELISA. **a)** Disaccharide composition of bioengineered rHS chains and percent site-specific disaccharide sulfation. **b)** Relative surface densities of rHS neoPGs after immobilization (100 ng/well) were determined via biotin-azide streptavidin-HRP assay. **c)** FGF1, FGF2, BMP2, BMP4, and VEGFA binding to rHS neoPGs immobilized at 100 ng/well. **d)** Ternary complex formation between FGF1 and FGFR1 or 2 in the presence of immobilized **Hep**_**7**_**-BSA** (100 ng/well) **e)** Effects of HSulf-1 processing of immobilized **Hep**_**7**_**-BSA** on FGF2, FGF1, or VEGFA binding (left). Differential processing of rHS neoPGs by HSulf-1 was assessed by FGF1 binding (right). (Bar graphs represent n = 3 replicates, *p*-values were determined using student’s t-test, \*\**p* < 0.01, \*\*\**p*< 0.001, \*\*\*\**p*< 0.001,).

Surface density of GAG chains determines the avidity of their protein interactions and can influence binding analysis. Hence, we set to determine whether the immobilization efficiency of neoPGs was impacted by the structure and valency of their pendant rHS chains. First, we quantified the surface density of unconjugated BSA and **Hep**_**7**_**-BSA** by taking advantage of the remaining unreacted cyclooctynes present on the protein. The molecules were adsorbed on plates (100 ng/well) in the presence of biotin-PEG_11_-azide to allow for quantification of their surface density via streptavidin-HRP ELISA (Figure 4b). Treating **BCN-BSA** and **Hep**_**7**_**-BSA** with increasing amounts of the biotinylating agent, we determined that equimolar quantity of biotin-PEG_11_-azide per total cyclooctyne in **BCN-BSA** (i.e., 16 equiv.) provided maximal labeling of both conjugates after immobilization (Figure S8). Analysis of all neoPGs using half the maximal amount of biotin-PEG_11_-azide to account for rHS attachment and provide equal labeling of all neoPG conjugates, showed uniform surface immobilization regardless of glycosylation status (Figure 4b). The uniform neoPG adsorption on the plate surface thus allows for direct comparison of their protein binding activities.

Growth factors and morphogens commonly engage GAGs and many are well known HS-binding proteins, though the structural basis for their recognition of HS remains elusive. We assessed the binding activity of the rHS neoPGs for established HS binding proteins (Figure 4c), including fibroblast growth factors 1 and 2 (FGF1, FGF2), bone morphogenic proteins 2 and 4 (BMP2, BMP4), and vascular endothelial growth factor a (VEGFA). It is known that FGF1 and FGF2 require 2-*O* sulfate on the iduronic acid (IdoA2S) residues in HS for binding.^38^ Accordingly, we observed preference for FGF1 and FGF2 binding to **rHS01**_**6**_**-BSA** (containing IdoA2S) over **rHS02**_**10**_**-BSA** (lacking 2-*O*-sulfates). The higher valency and overall charge of **rHS02**_**10**_**-BSA** did not compensate for the lack of 2-*O* sulfation.

We observed similar BMP2 binding profiles to what has been identified in current literature.^39^ Structures with reduced *N*-sulfation (**rHS01**_**6**_**-BSA**) present significantly diminished binding, which may be compensated for by increasing the rHS chain length and neoPG valency (**rHS02**_**10**_**-BSA**, ∼ 40 kDa for rHS02 vs ∼20 kDa vs for rHS01). While reduced sulfation levels in **rHS01**_**6**_**-BSA** and **rHS02**_**10**_**-BSA** reduced BMP4 binding, the presence of 3-*O*-sulfation in rHS08 restored BMP4 binding. For VEGFA, literature reports indicate a preference for disaccharides composed of iduronic acid and displaying 6-*O-*sulfation and *N*-sulfation with no consensus on the requirement for 2-*O*-sulfation or 3-*O*-sulfation.^40^ We observed strongest binding of VEGFA to **rHS08**_**6**_**-BSA** and weakest binding to **rHS01**_**6**_**-BSA** and **rHS02**_**10**_**-BSA**, which are similar based on quantitative levels of sulfation apart from the presence of 3-*O*-sulfation in **rHS08**_**6**_**-BSA**. For neoPGs carrying rHS structures with high levels of *N-* and 6-*O*-sulfation, **rHS09**_**10**_**-BSA** and **rHS29**_**3**_**-BSA**, a reduction in 2-*O*-sulfation and chain valency in the latter significantly decreased VEGFA binding.

Cell surface HS can also facilitate functional pairing of GFs with their cognate receptors. To assess the ability of the immobilized neoPGs to promote the formation of ternary GAG-GF-receptor complexes, we incubated recombinantly expressed ectodomains of FGF receptors FGFR-1 and -2 fused to the Fc domain of IgG with immobilized **Hep**_**7**_**-BSA** neoPG in the presence or absence of FGF1 (Figure 4d). FGFR-1 and -2 binding, detected using an anti-Fc antibody, only occurred in the presence of FGF1, indicating the formation of a ternary complex.

The neoPG array platform can also be used to analyze the interactions of GAG-modifying enzymes, such as the extracellular human 6-*O*-endosulfatase 1 (HSulf-1), with their substrates. HSulf-1 selectively cleaves 6-*O*-sulfates on cell-surface and ECM HSPGs and alters their GF binding activity;^41, 42, 43, 44^ however, the substrate specificity of this enzyme is still poorly defined.^45^

First, we tested the effects of HSulf-1 desulfation of immobilized **Hep**_**7**_**-BSA** on FGF1, FGF2, or VEGFA binding, which were selected based on their differential sensitivity to HSulf-1 activity.^46^ We observed significant loss of FGF1 and VEGFA binding but no significant change for FGF2 (Figure 4e), matching their known HS-binding dependence and independence on 6-*O*-sulfation of HS.^38,40^ Next, we investigated the effect of rHS composition on HSulf-1 activity and FGF1 binding (Figure 4e). We observed little change in FGF1 binding after HSulf-1 processing of rHS01 and rHS02 in their respective neoPGs, consistent with their overall low level of 6-*O*-sulfation. Notably, conjugate **rHS08**_**6**_**-BSA** which also features low overall 6-*O*-sulfation still exhibits reduced FGF1 binding. rHS08 specifically contains the 3-*O*-sulfation motif, which engendered the strongest binding by FGF1, and the strongest response to HSulf-1 activity. The neoPG reagents thus provide suitable probes for characterizing the substrate specificity of the HSulfs and their dynamic regulation HS interactions with signaling proteins.

### Activity of neoPGs in regulation of GF signaling

Correlating the interactions of HS structures with GFs observed in binding assays with their biological activities in cells is critical for establishing their structure-activity relationships. We assessed the ability of neoPGs to promote functional pairing between FGF2 and its cell surface receptor FGFR (Figure 5a). Cultured wild-type murine embryonic stem cells (WT-mESC) and HS-deficient mESCs, lacking the HS polymerizing enzyme, exostosine 1 (Ext1^-/-^), were serum starved and incubated with soluble **Hep**_**7**_**-BSA** or **Hep** (0.25 - 5 µg/mL). The mESCs were stimulated with and without soluble FGF2 (25 ng/mL) and probed for ERK phosphorylation (Phos-ERK) via Western blot analysis as a readout for FGFR activation and MAPK signaling (Figure 5b). **Hep**_**7**_**-BSA** enhanced FGF2-mediated FGFR signaling compared to equivalent concentrations of **Hep**, inferring an increased capacity of the multivalent neoPG to form a stable ternary complex compared to Hep alone. We also tested the impact of valency using **Hep**_**x**_**-BSA** (x = 1, 2, or 4, normalized to total Hep concentration) and observed increased ERK phosphorylation response with increasing conjugate valency (Figure 5c), consistent with a valency-dependent FGF2 binding observed for immobilized **Hep**_**7**_**-BSA** (Figure 3b). These findings demonstrate the importance of the overall macromolecular architecture of the neoPGs on signaling and the ability to correlate binding data obtained from ELISA assays to biological activity in cells.

**Figure 5.**
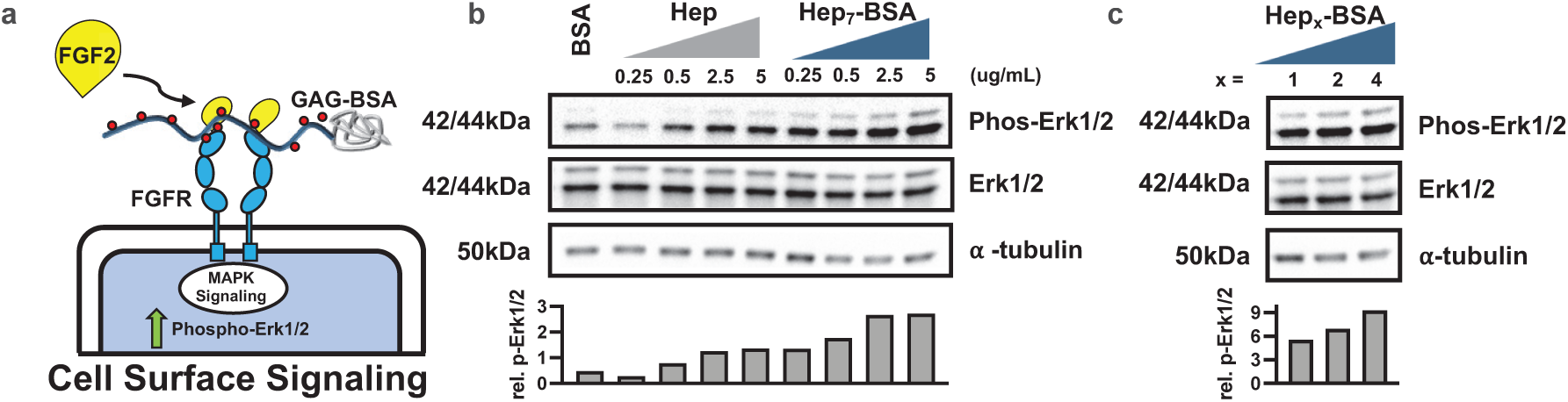
neoPGs promote FGF2 signaling in HS deficient mouse embryonic stem cells (Ext1^-/-^ mESCs). **a)** Cartoon depiction of FGF2 stimulation assay in Ext1^-/-^ mESCs in the presence of **Hep**_**x**_**-BSA. b)** The cells were stimulated with FGF2 (25 ng/mL) in the presence of increasing concentrations of soluble Hep or **Hep**_**7**_**-BSA** conjugate. Cell lysates were assesed for Erk1/2 phosphorylation after stimulation via western blot and densitometry analysis. α-Tubulin was used as a loading control. The data are representative examples of at least 2 independent biological replicates. **c)** Ext^-/-^ mESCs were stimulated with FGF2 (25 ng/mL) in the presence of **Hep**_**x**_**-BSA** with increasing numbers of Hep chains (x = 1, 2, 4) normalized to 5 µg/mL heparin concentration to assess valency effects on MAPK activation.

## Conclusion

In this report, we outline an efficient and tunable method for generating neoPG conjugates by merging biologically-derived or bioengineered GAG polysaccharides and cyclooctyne-modified BSA protein via a novel bifunctional fluorogenic linker. The method was applicable to all members of the GAG family, including rHS polysaccharides produced in cells with genetically engineered HS biosynthesis. The conjugation process generates a fluorescent signal, which can be used to monitor the progress of the reaction and determine the overall composition of the resulting neoPGs. The reagents were arrayed in 96-well plates to evaluate the ligand specificity of GAG-binding proteins in a convenient ELISA format. Compared to traditional immobilization strategies based on electrostatic adsorption, the conjugation of GAGs to BSA via their reducing-ends provides impartial surface presentation, which enabled analysis of substrate specificity for GAG-remodeling lyase and sulfatase enzymes. The neoPGs can also be deployed as soluble probes to confirm their growth factor binding activity in cell signaling assays. With the rapid advancements in precision engineering of cellular glycosylation pathways, these reagents are poised to provide a powerful complement to existing array platforms based on synthetically defined oligosaccharides and contribute a more complete understanding of structure-activity relationships in GAG-protein binding interactions.

## Methods and Materials

### 3-azido-coumarin-7-sulfonyl fluoride (ACS-F) synthesis

To a 4 mL vial containing 3-azido-7-hydroxycoumarin (100 mg, 0.49 mmol, 1.0 eq.) and 4-[(Acetylamino)phenyl]imidodisulfuryl difluoride (AISF, 186 mg, 0.58 mmol, 1.2 equiv.) was added anhydrous DMSO (1.6 mL) followed by 1,8-diazabicyclo [5.4.0]undec-7-ene (DBU, 161 µL, 1.08 mmol, 2.2 equiv.) over a period of 60 seconds. The reaction mixture stirred at ambient temperature for 1 h, diluted with ethyl acetate and washed with 0.5 N HCl (2x) and once with brine. Combined organic fraction was dried with anhydrous magnesium sulfate and concentrated under reduced pressure. The crude residue was purified by silica gel flash chromatography (0→ 40% EtOAc/ Hex with elution at ∼15% EtOAc/Hex) to afford the product (35 mg, 28% yield) as a crystalline clear solid. ^1^H NMR (500 MHz, CDCl_3_) δ 7.54 (d, *J* = 8.6 Hz, 1H), 7.37 (d, *J* = 2.3 Hz, 1H), 7.30 (ddd, *J* = 8.6, 2.4, 0.7 Hz, 1H), 7.21 (s, 1H). ^13^C NMR (126 MHz, CDCl_3_) δ 156.47, 151.52, 149.88, 128.93, 127.91, 124.00, 119.78, 118.10, 110.09 ppm. ^19^F NMR: (282 MHz, CDCl_3_) δ 39.0 (s, 1F). Absorbance and fluorescence spectra were obtained by analyzing unreacted and Heparin-conjugated to ACS-F by absorbance scan pedestal analysis on a Nanodrop 2000c or fluorescence excitation and emission scan on a fluorimeter, respectively.

### GAG conjugation to ACS-F

Commercial GAGs including heparin (20 mg, Iduron, Macclesfield SK10 4TG, UK), TEGA recombinant HS (rHS), or biologically-sourced GAGs were transferred to a PCR tube and dissolved in 90 µL of 1 M urea, 1 M sodium acetate, pH 4.5 buffer. To this solution was added 10 µL of a 1.15 M n-methylaminooxy-propylamine linker^34^ (11.5 µmoles). Reducing end conjugation proceeded at 50°C for 24-48 h. The reaction was quenched with 200 µL of 2 M Tris-HCl, pH 8.1 and GAG-amine was purified by PD-10 column, followed by concentration and removal of excess linker using 3 kDa molecular weight spin filters as per manufacturer’s instructions (Amicon, Millipore Sigma, St. Louis, MO). To 400 µL of recovered GAG-amine was added 200 µL of 100 mM sodium phosphate, pH 8.0 and 600 µL DMSO. ACSF (38 mg, 133 µmoles) was dissolved in 400 µL DMSO and transferred to the GAG-amine solution. Sulfonyl fluoride exchange (SuFEx) proceeded at ambient temperature, shaking for 24 h. The reaction was diluted with 900 µL water and similarly purified over PD-10 column with elutions collected in a CoStar clear 96-well plate and analyzed by microplate absorbance at 326 nm to visualize ACS-GAG and excess ACS-F fractions. ACS-GAG fractions were pooled and similarly concentrated by 3 kDa spin filtration, followed by lyophilization.

### BCN-BSA synthesis

To a 1.5 mL microcentrifuge tube (Fisher scientific, Cat. No. 05408129) was added 1 mL of 100 mM sodium phosphate buffer, pH 8.0, 10 mg BSA (VWR, Cat. No. 0332-25G), and 78.1 µL of a 10 mg/mL (1R,8S,9s)-Bicyclo[6.1.0]non-4-yn-9-ylmethyl N-succinimidyl carbonate (BCN, 17 eq.) (Sigma Aldrich, Cat. No. 744867-10MG) and stirred at 4°C for 16 h. The reaction was dialyzed against MilliQ water in 25 kDa molecular weight cut-off dialysis tubing (Spectra, Cat No. 132126), for 48 hours, replacing water after 24 hours. Lyophilization of the dialyzed product affords 11 mg of the product (quantitative yield). MALDI-TOF MS analysis indicates the modified BSA protein has a molecular weight of about 69,689 daltons compared a starting mass of 66,808 daltons for unmodified BSA. Each additional BCN adds 177.3 daltons, a difference of 3,259 daltons indicates approximately 16 BCN/BSA.

### ACS-GAG conjugation to BCN-BSA

To the wells of a black, clear-bottom CoStar 96-well plate were added either 200 µL PBS, 50 µL of 1 mM ACS-F (50 nmoles), 50 µL of 1 mM ACS-GAG (50 nmoles), 100 µL BCN-BSA (0.5 mg/mL in water, 0.9 nmoles BSA and 8 nmoles BCN), 50 µL ACS-F + 100 µL BCN-BSA, or 50 µL ACS-GAG + 100 µL BCN-BSA. Each well was brought up to a final volume of 200 µL with PBS. Using a microplate spectrophotometer, kinetic fluorescence readings were collected with Ex. 393 nm/ Em. 477 nm at various time points initially after addition of all reagents to 24-48 h. Post-incubation at ambient temperature for 26 h, wells containing neoPG (GAG-BSA) or TCS-BSA (triazole) were filtered (5 × 500 µL water) through a 30 kDa molecular weight spin filter to remove unreacted GAG and ACS-F. The recovered 40 µL of neoPG or TCS-BSA was transferred into separate PCR tubes and water was added to a final concentration of 200 ug/mL BSA assuming 96% BSA recovery based on manufacturers data sheet and confirmed by BCA assay. The TCS-BSA and neoPG samples were further diluted for adsorption onto 96-well plates for binding assays.

### 96-well ELISA binding assays

neoPGs were diluted into 1X Dulbecco’s phosphate buffered saline w/ Ca^2+^ and Mg^2+^ (PBS) and adsorbed to a Greiner microplate high-binding clear-bottom 96-well plate (100 ng/well, 100 µL final volume per well). Conjugates were adsorbed for 16 h at 4°C. Post-adsorption, wells were washed three times with PBS, blocked with 100 µL per well of 2% BSA in PBS for 1 h at ambient temperature, and washed again three times with PBS. Growth factors (GFs) and antibodies were diluted to final concentrations in 1% BSA/PBS following manufacturer instructions or otherwise noted and added to the wells in triplicate. A triplicate set of control wells were not incubated with GAG-binding proteins. Binding proteins incubated for 2 h at ambient temperature. Wells were then washed three times with PBS, then incubated with a respective antibody precomplexed with an HRP-conjugated secondary antibody as necessary. For a precomplex, primary and secondary antibodies or binding proteins and their antibodies were combined in 1.5 mL Eppendorf tubes in 200 µL of 1% BSA/PBS for 30 mins on ice prior to diluting to the final calculated concentration. Wells were incubated with the antibody precomplex for 1.5 h at ambient temperature, washed three times with PBS, and visualized with 100 µL of TMB substrate (Invitrogen, 00-4201-56) with resulting colorimetric product measured on a SpectraMax i3x plate reader under a kinetic cycle with 370 nm absorbance (10 min total with 30 second increments).

For enzymatic treatments including heparinase, chondroitinase ABC, hyaluronidase, keratanase II, and 6-O-endosulfatase 1 (HSulf-1), the neoPG immobilized plates were similarly blocked and washed with PBS as above followed by incubation in reaction buffer with or without enzyme present at 37°C for 16 h. Post-enzymatic treatment was followed by three PBS washes and similar protein binding procedures as described above. For Sulf-1 enzyme purification, A375 KDM2B^-/-^ cells, kindly provided by Dr. Jeffrey Esko (UCSD), were cultured in OptiMEM for 3-5 days, conditioned media was collected and filtered through a 50 kDa cut-off filter, total protein quantified by BCA assay, and presence of HSulf-1 validated by Western blot analysis and anti-Sulf1 detection. No detectable heparin lyase activity was present in conditioned media.

For biotin-PEG_11_-azide analyses, BCN-BSA and Hep_7_-BSA were immobilized onto high-binding 96-well plates in the presence of varying equivalents of the biotin-azide reagent compared to total BCN per BSA (with ∼17 BCN/BSA, 0.1 eq = 1.7 biotin-PEG_11_-azide per BSA and 1 eq = 17 biotin-PEG_11_-azide per BSA). To quantify neoPG immobilization efficiency, 0.5 eq of biotin-PEG_11_-azide was utilized in the immobilization assay to account for reacted BCN molecules that were no longer available. Post-immobilization at 4°C for 16 h, wells were washed three times with PBS, blocked with 2% BSA/PBS, and incubated with HRP-conjugated streptavidin for 1.5 h at ambient temperature. Colorimetric analysis of streptavidin binding was conducted similarly to above binding analyses using a kinetic cycle of 370 nm absorbance readings.

### Cell surface FGF receptor stimulation assay

FGF2 stimulation and western blotting was performed as detailed previously by the Godula lab.^34^ Briefly, mouse embryonic stem cells with endogenous HS production knocked out (Ext1^-/-^) were cultured in 6-well plates treated with 0.1% gelatin before being serum starved for 20 h in mESC growth medium lacking FBS. Cells were then treated with 25 ng/mL recombinant human basic FGF (Peprotech) in serum-free medium with heparin, BSA, or Hep_7_-BSA conjugates (5 µg/mL) for a duration of 15 min at 37°C, 5% CO2. The cells were then immediately chilled and lysed using a 1X RIPA lysis buffer supplemented with PMSF (1 mM) and 1X protease/phosphatase inhibitor cocktail. Cell lysates were analyzed by BCA assay to determine total protein concentration. 10 µg total protein from each sample was resolved on a 10% SDS-PAGE gel and transferred to a PVDF membrane for blotting. The membrane was blocked with 5% BSA in TBS supplemented with 0.1% Tween-20 (TBST) for a minimum of 1 h at ambient temperature prior to staining (overnight, 4°C) with anti-phospho Erk, anti-total Erk, or anti-alpha tubulin (1:1250, 1:1250, 1:25000 respectively in 5% BSA). After three TBST washes the membrane was incubated with HRP-conjugated secondary antibodies (1:2,000 anti-rabbit HRP and 1:10000 anti-mouse HRP, respectively) for approximately 1.5 h at ambient temperature.

Following a series of TBST washes the blots were visualized using Luminata Forte HRP detection reagent and imaged on a gel scanner (BioRad) for chemiluminescence. For sequential staining, blots were washed in TBST, stripped using Restore PLUS Western blot stripping buffer, washed again in TBST and blocked in 5% BSA for at least 1 h at ambient temperature before further staining. Images were analyzed using ImageJ, with phospho-Erk1/2 and total-Erk1/2 normalized to alpha tubulin, then phospho-Erk was normalized to total-Erk. Lastly, the levels of relative Erk phosphorylation were determined by setting the phosphorylation of Erk in samples containg Ext1^-/-^ mESCs without FGF2 or neoPG to equal 1.

## Acknowledgements

We thank Dr Jeffrey D. Esko (UC San Diego) for helpful discussions and for generously providing reagents. This work was supported by the NHLBI K12 Career Development in Glycosciences program K12HL141956 to R.N.P. This work was supported in part by the NIH Director’s New Innovator Award (NICHD: 1DP2HD087954-01). KG was supported by the Alfred P. Sloan Foundation (FG-2017-9094) and the Research Corporation for Science Advancement via the Cottrell Scholar Award (grant #24119).

## Author Contributions

R.N.P., K.G., and P.L.S.M.G. conceived, initiated, and coordinated the project. R.N.P., J.L.F, S.P., B.T., D.K., and L.L. designed and performed the experimental work. B.E.T. and C.A.G. provided rHS samples from TEGA Therapeutics. R.N.P., J.L.F., and K.G. wrote the manuscript. All authors discussed the experiments and results, read, and approved the manuscript.

