## Supplemental for "Biologically-derived neoproteoglycans for profiling protein-glycosaminoglycan interactions"

**Table of Contents:**

Abbreviations ....................................................................................................................S4

**Table S1.** Biological reagents and consumables .................................................. S5-6

Materials and Instrumentation .......................................................................................S7

**A. Synthesis of 3-azidocoumarin-7-sulfonyl fluoride (ACS-F)** ......................... S7-S10

General chemical synthesis of **ACS-F** ..........................................................................S7

**Figure S1.** $^{1}$H NMR of **ACS-F** ..........................................................................S8

**Figure S2.** $^{13}$C NMR of **ACS-F** ..........................................................................S9

**Figure S3.** $^{19}$F NMR of **ACS-F** ..........................................................................S10

**Figure S4.** IR Spectra of **ACS-F** ...............................................................................S10

**B. Preparation and characterization of BCN-BSA** ....................................................S11

General chemical synthesis of **BCN-BSA** ......................................................................S11

**Figure S5.** MALDI-TOF spectra of **BCN-BSA** ...........................................................S11

**C. Preparation and characterization of neoPG conjugates** ...............................S12-16

Aminooxy-amine conjugation to Hep and TAMRA validation ..........................................S12

Synthesis of Hep-amine ...................................................................................................S12

Synthesis of Hep-TAMRA .................................................................................................S12

**Figure S6.** Hep-TAMRA PD-10 column elution profile ...............................................S12

Heparin-amine conjugation to **ACS-F** via SuFEx chemistry ..........................................S13

Synthesis of **Hep-ACS** ......................................................................................................S13

**Figure S7. Hep-ACS** PD-10 column elution profile ....................................................S13

**ACS-F** conjugation to **BCN-BSA** .........................................................................................S14

Synthesis of **TCS-BSA** via SPAAC ....................................................................................S14

**Figure S8.** MALDI-TOF spectra of **TCS-BSA** ............................................................S14

Standard curve of **TCS-BSA** fluorescence .......................................................................S14

**Figure S9. TCS-BSA** fluorescence standard curve ....................................................S15

Determination of heparin concentration via the carbazole assay ...................................S15

Determination of BSA concentration via the BCA assay .................................................S15

**Figure S10.** Analysis of **Hep-BSA** composition via carbazole and BCA assays ..S15

**Table S2.** Composition of neoPGs ...................................................................................S16

NeoPG Structures and Conjugation Valency ....................................................................S16

**D. Binding assays and functional assays with HS-BSA conjugates** ................S17-19

Biotin-azide immobilization assay .........................................................................S17

**Figure S11.** Biotin labeling of immobilized **BCN-BSA** and **Hep-BSA** ...................S17

**Figure S12.** Dose-dependent FGF1 binding to rHS-BSA conjugates ....................S18

**Figure S13.** Dose-dependent FGF2 binding to rHS-BSA conjugates ....................S18

**Figure S14.** FGF2 stimulation of Ext $1^{-/-}$ mESCs in presence of Hep and **Hep$_7$-BSA** ..............................................................................................................S19

**Figure S15.** FGF2 stimulation of Ext $1^{-/-}$ mESCs in presence of **Hep$_X$-BSA** (x = 1, 2, 4) ..............................................................................................................S19

**References** ..........................................................................................................S20

### ABBREVIATIONS

**ACS-F =** 3-azidocoumarin 7-sulfonyl fluoride

**BCA =** bicinchoninic acid

**BCN =** bicyclo[6.1.0]nonyne

**BMP =** bone morphogenic protein

**BSA =** bovine serum albumin

**DCM =** dichloromethane

**DMSO =** dimethyl sulfoxide

**ELISA =** enzyme-linked immunosorbent assay

**ESI-MS =** electrospray ionization mass spectrometry

**EtOAc =** ethyl acetate

**FGF =** fibroblast growth factor

**HCl =** hydrochloric acid

**Hep =** heparin

**Hex =** n-hexanes

**HRP =** horse radish peroxidase

**HS =** heparan sulfate

**IR =** infrared spectroscopy

**MALDI-TOF =** matrix-assisted laser desorption/ionization – time of flight

**MQ =** milli-Q ultrapure water

**MWCO =** molecular weight cut-off

**NHS =** N-hydroxysuccinimide

**NMR =** nuclear magnetic resonance

**PBS =** phosphate buffered saline

**PD-10 =** prepacked sephadex® G-25 medium disposable column

**RBF =** round-bottom flask

**SuFEx =** Sulfur (IV) fluoride exchange

**TCS** = triazole coumarin sulfonyl

**TFA =** trifluoroacetic acid

**VEGF =** vascular endothelial growth factor

**TABLE S1. REAGENTS AND CONSUMABLES**

| Biological Reagents | Source | Catalog No. |
|---|---|---|
| Anti-alpha tubulin | Cell Signaling | 3873s |
| Anti-BMP2 | R&D Systems | MAB3551 |
| Anti-BMP4 | Peprotech | 500-M121-500UG |
| Anti-Erk1/2 | Cell Signaling | 4695s |
| Anti-FGF1 | Invitrogen | PA5-79249 |
| Anti-FGF2 | Millipore | 05-118 |
| Anti-mouse IgG HRP | Cell Signaling | 7076s |
| Anti-mouse light chain specific HRP | Millipore | AP-200P |
| Anti-pErk1/2 | Cell Signaling | 4370s |
| Anti-rabbit IgG HRP | Cell Signaling | 7074S |
| Anti-VEGF | Invitrogen | P802 |
| BMP2 | R&D Systems | 355-BM |
| BMP4 | Peprotech | 120-05 |
| Bovine Serum Albumin (BSA) | Spectrum | A3611 |
| CD44-Fc | Biolegend | 783804 |
| Chondroitin Sulfate from Bovine Cartilage | Sigma | C6737-5G |
| Chondroitinase ABC | Sigma | C3667 |
| CS-56 Anti-CS | Sigma | C8035 |
| Fibroblast growth factor 1 | Abcam | Ab91374 |
| Fibroblast growth factor 2 | Peprotech | 100-18B |
| Fibroblast growth factor receptor 1α | Abcam | Ab55758 |
| Fibroblast growth factor receptor 2α (IIIc) | R&D Systems | 712-Fr |
| Goat anti-human IgG HRP | Invitrogen | 31418 |
| Heparin | Iduron | HEP001 |
| Heparinase I, II, III recombinantly purified | Gift from Jeffrey Esko lab | |
| Hyaluronan LMW 40,000-50,000 | Carbosynth | FH01773 |
| Rabbit anti-FGF2 | Sigma Aldrich | 05-118 |
| Recombinant HS (01, 02, 08, 09, 29) | TEGA Therapeutics | rHS-01,02,08,09,29 |
| Streptavidin-HRP | Raybiotech | EL-HRP |
| Vascular endothelial growth factor a | Thermo | PHC9394 |

| | | |
|---|---|---|
| Vascular endothelial growth factor receptor 1 | R&D Systems | 321-FL |
| Vascular endothelial growth factor receptor 2 | R&D Systems | 357-KD |
| **Chemical Reagents** | **Source** | **Catalog No.** |
| 1,8-diazabicyclo [5.4.0]undec-7-ene (DBU) | Sigma Aldrich | 139009-25G |
| 2-aminoacridone | Sigma Aldrich | 06627 |
| 3-azido-7-hydroxycoumarin | Carbosynth | FA31762 |
| 4-(Acetylamino)phenyl]imidodisulfuryl difluoride (AISF) | Sigma Aldrich | 901243 |
| Biotin-dPEG$_{11}$-Azide | Quanta Biodesign | 10784 |
| Carbazole | Ultra Scientific | HAH-022 |
| NHS-BCN | Sigma Aldrich | 744867-10MG |
| TAMRA-NHS | Invitrogen | C300 |
| Triton X-100 | Alfa Aesar | A16046 |
| Tween-20 | VWR | M147-4L |
| **Consumables** | **Source** | **Catalog No.** |
| 3 kDa Centrifugal Spin Filters | Amicon | UFC500396 |
| 50 kDa Centrifugal Spin Filters | Amicon | UFC505096 |
| 25 kDa Dialysis Tubing | Spectra | 132126 |
| 1.5 mL Microcentrifuge Tube | Thermo Fisher | 5408129 |
| Microtiter Plate Black clear-bottom 96-well | Corning | 3915 |
| Microtiter Plate Clear 96-well | Corning | 3370 |
| Microtiter Plate EIA/RIA half-area high-binding 96-well | Corning | 3690 |
| Microtiter Plate Black clear-bottom high-binding 96-well | Greiner | 655097 |
| PCR tube | Thermo Fisher | AB0337 |
| PD-10 Desalting Column | GE Life Sciences | 17085101 |
| Phosphate Buffered Saline with Ca/Mg | Corning | 21-030-CM |
| TMB substrate solution | Invitrogen | 00-4201-56 |

### MATERIALS AND INSTRUMENTATION

All chemical and biological reagents and solvents were sourced as indicated in Table S1 and used as received. 3-((methylamino)oxy)propan-1-amine (n-methylamin was synthesized as previously reported.[1] Nuclear magnetic resonance (NMR) spectra were collected on a Bruker 300MHz NMR spectrometer. Spectra are reported in parts per million (ppm) on the δ scale relative to the residual solvent as an internal standard. Data are reported as follows: chemical shift (s = singlet, d = doublet, dd = doublet of doublets, t = triplet, q = quartet, br = broad, m = multiplet), coupling constants (Hz), and integration. Absorbance and fluorescence values for 96-well plate applications were collected on a SpectraMax i3x plate reader (Molecular Devices) with SoftMax Pro software. Microtiter plate analysis was conducted on a SpectraMax C3 plate reader. All data were plotted and analyzed on GraphPad Prism 8 software. Fluorescence excitation and emission scanned on a Horiba FluoroMax4 Spectrofluorometer.

#### A. Synthesis of 3-azido-7-sulfonyl coumarin fluoride (ACS-F)

**Chemical synthesis of ACS-F:** To a 4 mL vial containing 3-azido-7-hydroxycoumarin (100 mg, 0.49 mmol, 1.0 eq.) and 4-(Acetylamino)phenyl]imidodisulfuryl difluoride (AISF, 186 mg, 0.58 mmol, 1.2 equiv.) was added anhydrous DMSO (1.6 mL) followed by 1,8-diazabicyclo [5.4.0]undec-7-ene (DBU, 161 µL, 1.08 mmol, 2.2 equiv.) over a period of 60 seconds. The reaction mixture stirred at ambient temperature for 1 h, diluted with ethyl acetate and washed with 0.5 N HCl (2x) and once with brine. Combined organic fraction was dried with anhydrous magnesium sulfate and concentrated under reduced pressure. The crude residue was purified by silica gel flash chromatography (0 $\rightarrow$ 40% EtOAc/ Hex with elution at ~15% EtOAc / Hex) to afford the product (35 mg, 28% yield) as a crystalline clear solid. $^{1}$H NMR (500 MHz, $CDCl_3$) δ 7.54 (d, *J* = 8.6 Hz, 1H), 7.37 (d, *J* = 2.3 Hz, 1H), 7.30 (ddd, *J* = 8.6, 2.4, 0.7 Hz, 1H), 7.21 (s, 1H). $^{13}$C NMR (126 MHz, $CDCl_3$) δ 156.47, 151.52, 149.88, 128.93, 127.91, 124.00, 119.78, 118.10, 110.09 ppm. $^{19}$F NMR: (282 MHz, $CDCl_3$) δ 39.0 (s, 1F). Absorbance and fluorescence spectra were obtained by analyzing unreacted and Heparin-conjugated to ACSF by absorbance scan by pedestal analysis on a Nanodrop 2000c or fluorescence excitation and emission scan on a fluorimeter.

**Figure S1.** $^1$H NMR of **ACS-F**

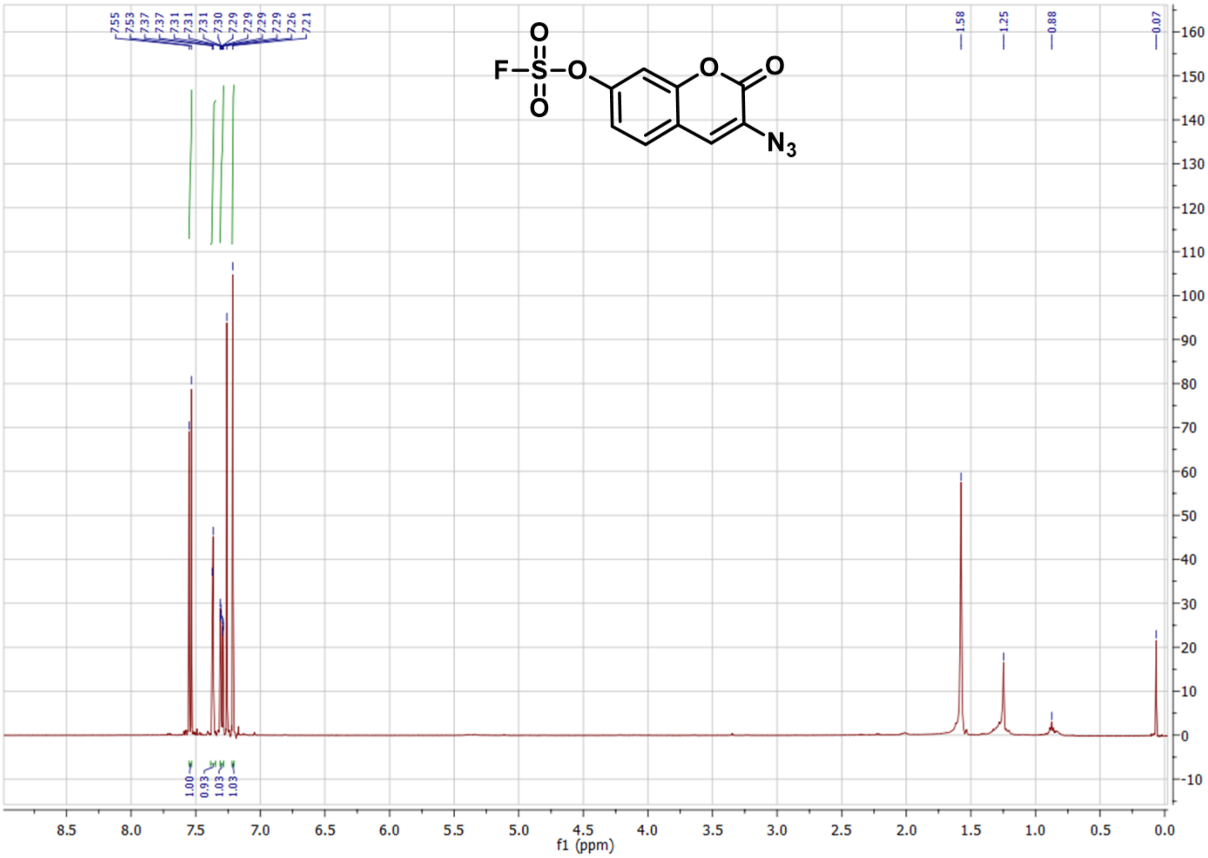

**Figure S2.** $^{13}$C NMR of **ACS-F**

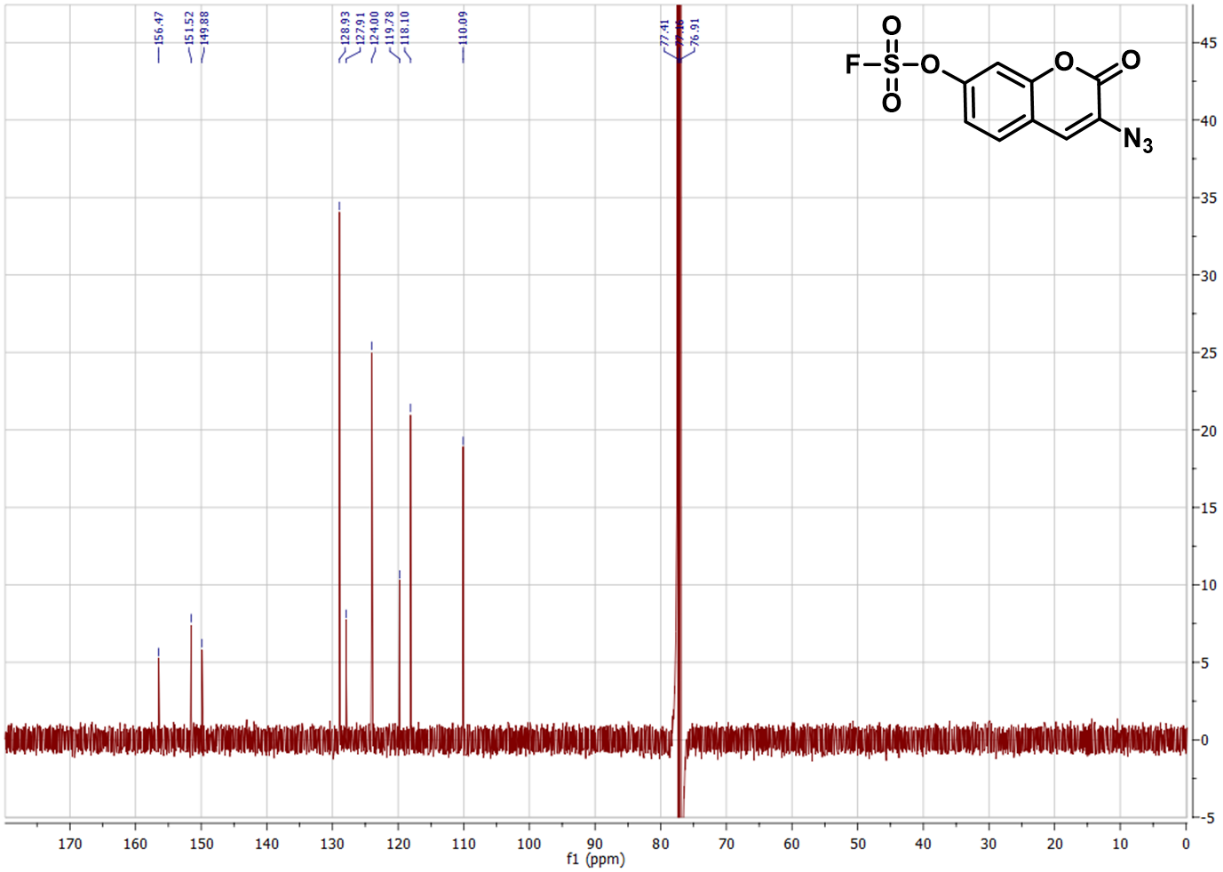

**Figure S3.** $^{19}$F NMR of **ACS-F**

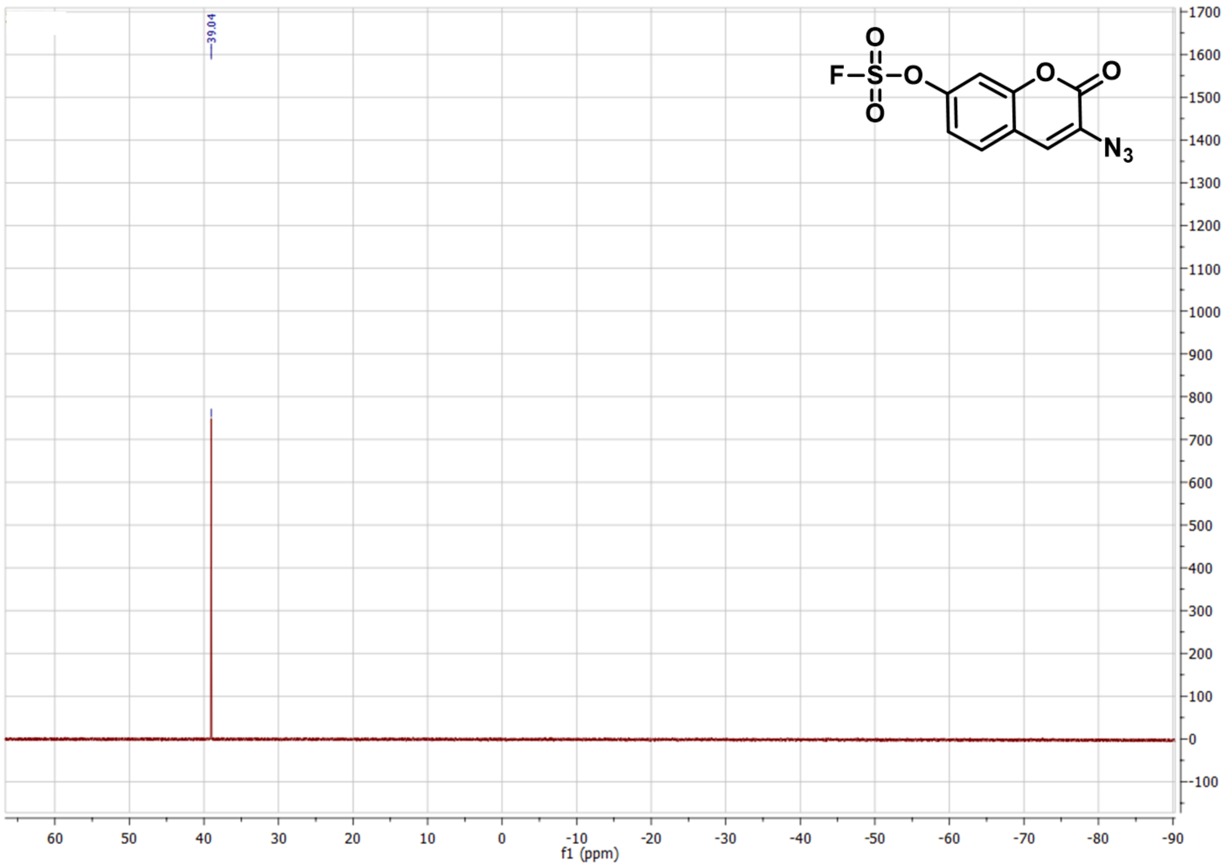

**Figure S4.** IR Spectra of **ACS-F**

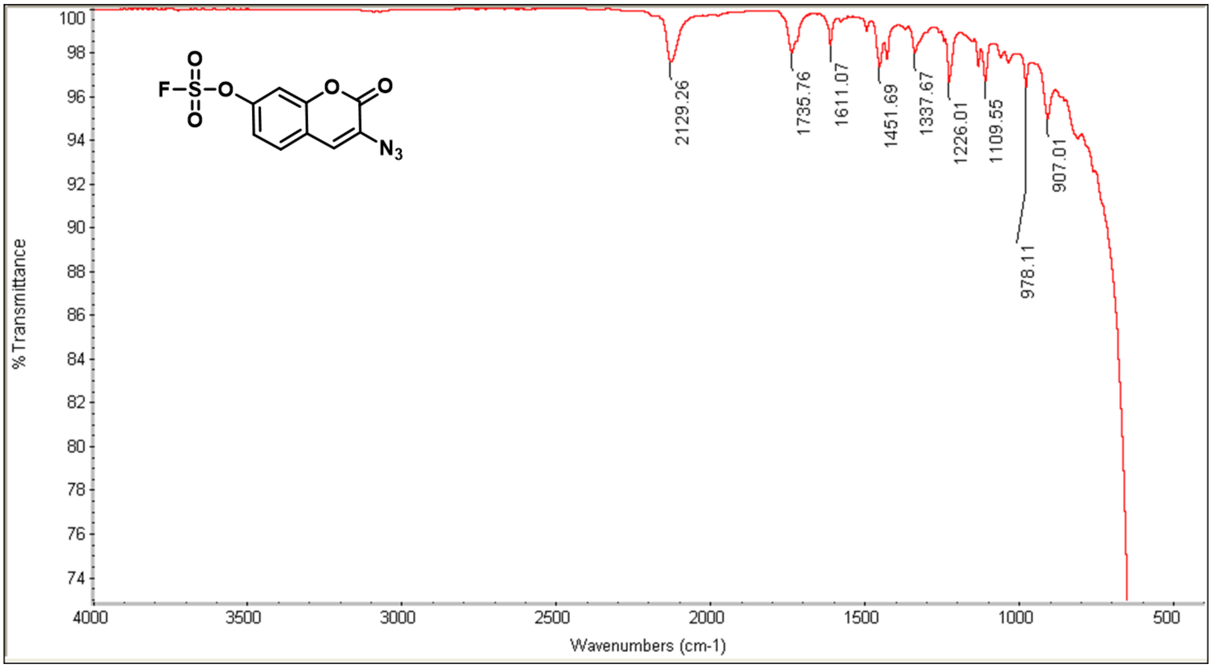

### B. Preparation and characterization of BCN-BSA

**Chemical synthesis of BCN-BSA:** To a 1.5 mL microcentrifuge tube (Fisher scientific, Cat. No. 05408129) was added 1 mL of 100 mM sodium phosphate buffer, pH 8.0, 10 mg BSA (VWR, Cat. No. 0332-25G), and 78.1 µL of a 10 mg/mL (1R,8S,9s)-Bicyclo[6.1.0]non-4-yn-9-ylmethyl N-succinimidyl carbonate (BCN, 17 eq.) (Sigma Aldrich, Cat. No. 744867-10MG) and stirred at 4°C for 16 h. The reaction was dialyzed against MilliQ water in 25 kDa molecular weight cut-off dialysis tubing (Spectra, Cat No. 132126), for 48 hours, replacing water after 24 hours. Lyophilization of the dialyzed product affords 11 mg of the product (quantitative yield). MALDI-TOF MS analysis indicates the modified BSA protein has a molecular weight of about 69,689 daltons compared to a starting mass of 66,808 daltons for unmodified BSA. Each additional BCN adds 177.3 daltons, a difference of 2,881 daltons indicates approximately 16 BCN/BSA.

**Figure S5.** MALDI-TOF spectra of **BCN-BSA**

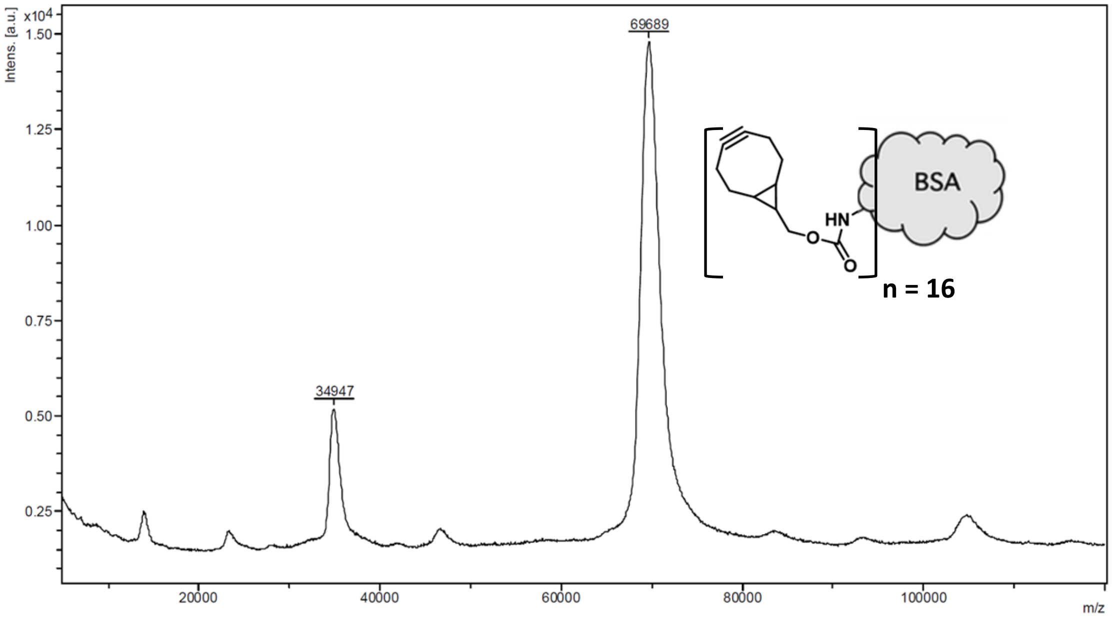

### C. Preparation and characterization of neoPG conjugates

#### Aminooxy-amine conjugation to Hep and TAMRA validation

**Synthesis of Hep-amine**: Heparin (20 mg) or TEGA recombinant HS (rHS) was added to a PCR tube and dissolved in 90 µL of 1 M Urea, 1 M sodium acetate, pH 4.5 buffer. To this solution was added 10 µL of a 1.15 M methylaminooxy-propylamine linker[1] (11.5 µmoles). Reaction proceeded at 50°C for 24 h. Reaction was quenched with 200 µL of 2 M Tris-HCl, pH 8.1 and heparin was purified by PD-10 column, followed by concentration and removal of excess linker using 3 kDa molecular weight spin filters.

**Synthesis of Hep-TAMRA:** To 10 µL of 20 mg/mL heparin-amine was added 40 µL of 100 mM sodium phosphate, pH 8.0 and 50 µL of 1 mM TAMRA-NHS. To a separate vial was added a comparable amount of aminooxy-amine linker to phosphate buffer and TAMRA-NHS as a control. After 24 h at ambient temperature with shaking, the two reactions were each separated by PD-10 column chromatography according to manufacture conditions, with water elution fractions collected in a black CoStar 96-well plate (~5 drops/well), and the fluorescence read on a SpectraMax plate reader at Ex. 550 nm/Em. 584 nm.

**Figure S6. Hep-TAMRA** PD-10 column elution profile

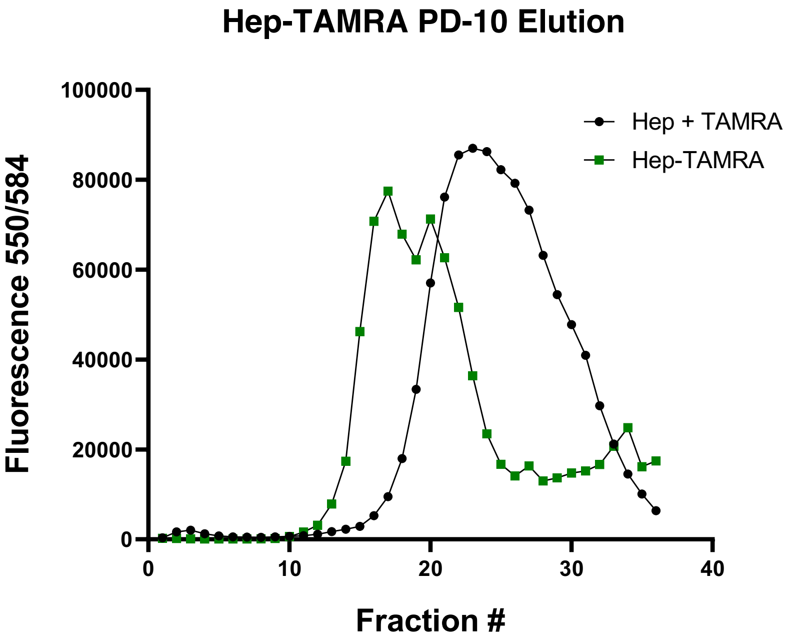

**Hep-amine conjugation to ACS-F via SuFEx chemistry**

**Synthesis of Hep-ACS:** To 400 µL of recovered heparin-linker was added 200 µL of 100 mM sodium phosphate, pH 8.0 and 600 µL DMSO. ACSF (38 mg, 133 µmoles) was dissolved in 400 µL DMSO and transferred to the heparin-linker solution. Conjugation proceeded at ambient temperature, stirring for 24 h. Reaction was diluted with 900 µL water and similarly isolated by PD-10 column with elutions collected in a CoStar clear 96-well plate and analyzed by microplate absorbance at 326 nm to visualize heparin-ACS and excess ACSF fractions. Heparin fractions were pooled and similarly concentrated by 3 kDa spin filtration, followed by lyophilization with ~53% recovery of tagged heparin.

**Figure S7. Hep-ACS** PD-10 column elution profile

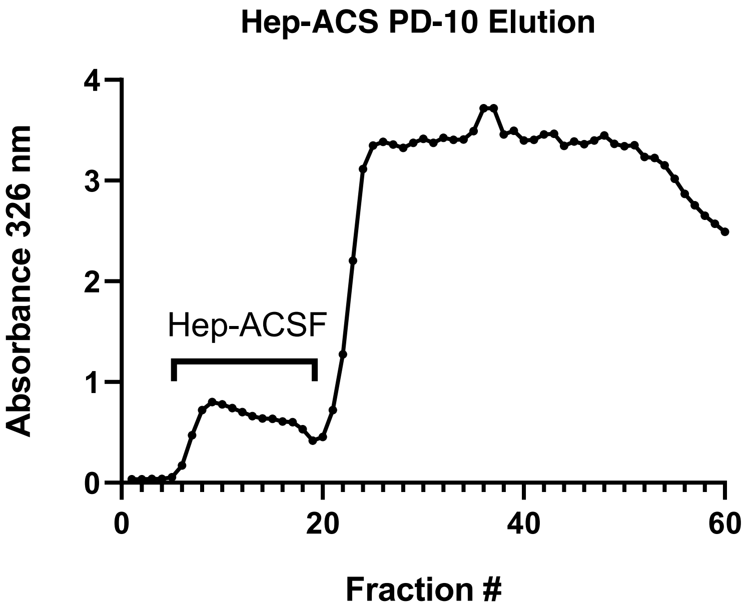

### ACS-F conjugation to BCN-BSA

**Synthesis of TCS-BSA via SPAAC:** To the wells of a black, clear-bottom CoStar 96-well plate were added either 200 µL PBS, 50 µL of 1 mM ACS-F (50 nmoles), 50 µL of 1 mM **Hep-ACS** (50 nmoles), 100 µL **BCN-BSA** (0.5 mg/mL in water, 0.9 nmoles BSA and 8 nmoles BCN), 50 µL **ACS-F** + 100 µL **BCN-BSA**, or 50 µL **Hep-ACS** + 100 µL **BCN-BSA**. Each well was brought up to a final volume of 200 µL with PBS. Using a microplate spectrophotometer, kinetic fluorescence readings were collected with Ex. 393 nm/ Em. 477 nm at various time points initially after 1 minute up to over 24 h. Post-incubation at ambient temperature for 26 h, wells containing BSA were filtered (5 x 500 µL water) through a 30 kDa molecular weight spin filter to remove excess heparin or **ACS-F**. Transferred the 40 µL recovered samples into individual PCR tubes and added water to final concentration of 200 ug/mL **BCN-BSA** assuming 96% **BCN-BSA** recovery based on manufacturer's spin filter manual. These BSA, **TCS-BSA**, and **Hep-BSA** samples were further diluted for binding studies. MALDI-TOF MS analysis indicates the modified **TCS-BSA** protein has a molecular weight of about 75,007 daltons compared to a starting mass of 66,808 daltons for BSA. Each additional TCS-BCN adds 477.1 daltons, a difference of 8,199 daltons indicates approximately 17 TCS.

**Figure S8.** MALDI-TOF spectra of **TCS-BSA**

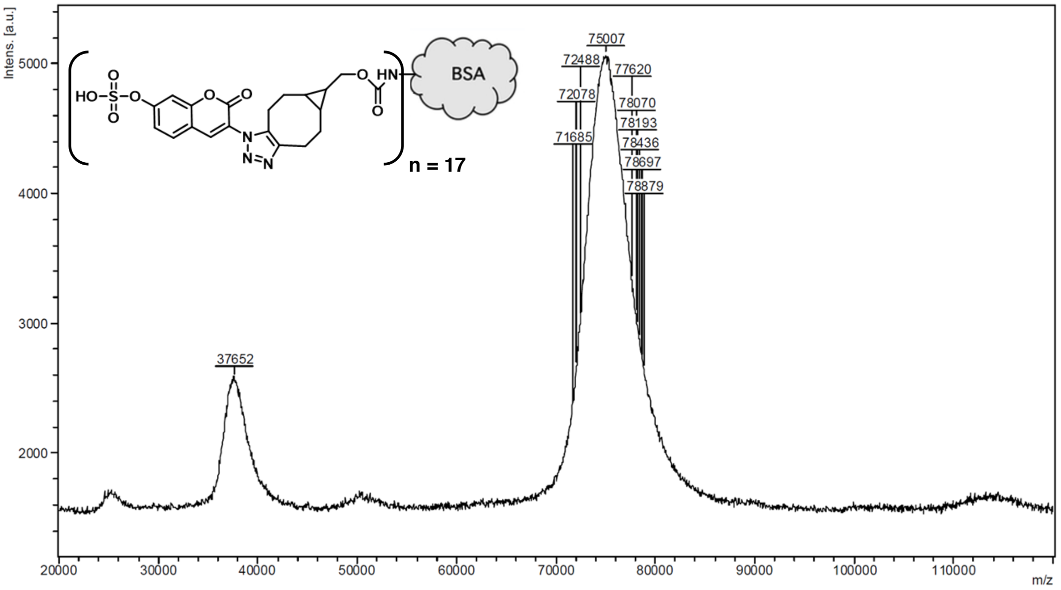

**Standard curve of TCS fluorescence:** In a glass reaction vial, combined 1 mM unconjugated BCN in DMF with 20 mM **ACS-F** in DMSO and let incubate at ambient temperature for 16 h stirring. Diluted this mixture into a dilution series in triplicate ranging from 10 nM to 100 µM BCN and measured fluorescence intensity (Ex. 393 nm/ Em. 477 nm) in a black 96-well plate. Similarly, **$HS_x$-BSA** fluorescence intensity was measured against the standard curve (Fig. S9) to approximate TCS concentration. Calculated TCS concentrations were combined with subsequent protein and GAG analyses to quantify the conjugation efficiency of HS to BSA protein, which was in accordance with approximate conjugation efficiencies from the ratio of **HS-BSA** end-fluorescence intensity to **TCS-BSA** during kinetic conjugation analysis.

**Figure S9. TCS** fluorescence standard curve

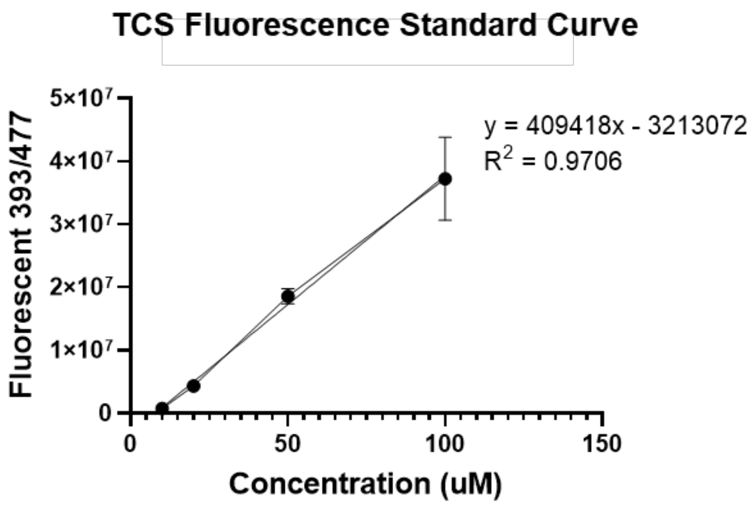

**Determination of heparin concentration via the carbazole assay:** This procedure was conducted as previously published and in accordance with standard protocols.[2]

**Determination of BSA concentration via the BCA assay:** This procedure was conducted in accordance with standard protocols.[3]

**Figure S10.** Analysis of **Hep-BSA** composition via carbazole and BCA assays

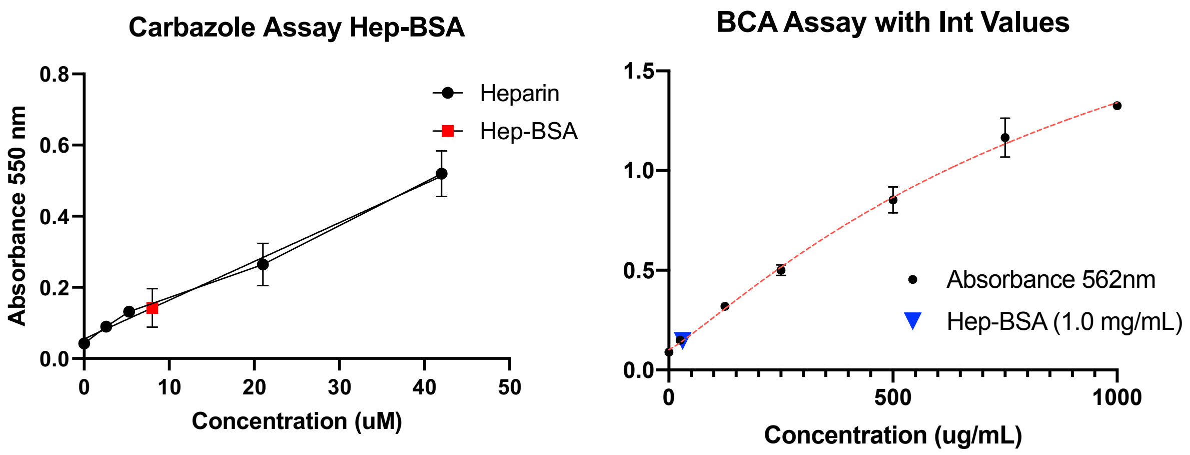

**Table S2.** neoPG structures and conjugation valency

| Commercial GAG | MW | Sulfation | BSA-Conjugation Valency |
|---|---|---|---|
| Heparin | 15 kDa | 95% | 7 |
| Bovine Cartilage CS | 30 kDa | - | 12 |
| LMW HA | 40—50 kDa | 0% | 6 |
| Recombinant GAG | MW | Sulfation | BSA-Conjugation |
| rHS01 | 20 kDa | 40% | 6 |
| rHS02 | 40 kDa | 50% | 10 |
| rHS08 | 15 kDa | 50% | 6 |
| rHS09 | 20 kDa | 97% | 10 |
| rHS29 | 18 kDa | 98% | 3 |
| Tissue Purified GAG | MW | Sulfation | BSA-Conjugation |
| Pig Lung HS | - | 64% | 8 |
| Pig Lung CS | - | - | 8 |
| Pig Lung KS | - | - | 6 |
| Mouse Liver HS | - | 12% | 6 |

**NeoPG structures and conjugation efficiencies.** Molecular weight (MW) of commercial GAGs were reported in provided data sheets. MW of rHS structures determined by size-exclusion chromatography analysis with standard curve by TEGA Therapeutics and reported in data sheets. Tissue purified GAGs not analyzed for MW. Sulfation was either provided by supplier or analyzed by disaccharide compositional analysis using high performance anion exchange chromatography as a service provided by UCSD Glycoanalytics Core. BSA-Conjugation efficiency calculated by ratio of fluorescence intensity of GAG-BSA to TCS-BSA at end-point of conjugation kinetic analysis and confirmed by TCS standard curve measurement in combination with carbazole and BCA assay data.

### D. Binding assays and functional assays with HS-BSA conjugates

**Biotin-azide immobilization assay:** Protocol described in Materials and Methods section of main text. Briefly, BCN-BSA and $Hep_7$-BSA (100 ng/well) were immobilized onto high-binding 96-well plates in the presence of biotin-$PEG_{11}$-azide reagent. Equivalents of biotin-azide were comparable to total BCN per BSA (with <17 BCN/BSA, 0.1 eq = 1.7 biotin-$PEG_{11}$-azide per BSA and 1 eq = 17 biotin-$PEG_{11}$-azide per BSA). In order to saturate BCN on the BSA, up to 25 equivalents (425 biotin-$PEG_{11}$-azide per BSA) were added. Biotin-$PEG_{11}$-azide equivalents were included without BSA present, serving as a background. Post-immobilization at 4°C for 16 h, wells were washed three times with PBS, blocked with 2% BSA/PBS, and incubated with HRP-conjugated streptavidin for 1.5 h at ambient temperature. Colorimetric analysis of streptavidin binding was conducted similarly to above binding analyses using a kinetic cycle of 370 nm absorbance readings. * = P-value <0.05, ** = P-value <0.01 using Prism statistical analysis student's t-test.

**Figure S11.** Biotin labeling of immobilized **BCN-BSA** and **Hep-BSA**

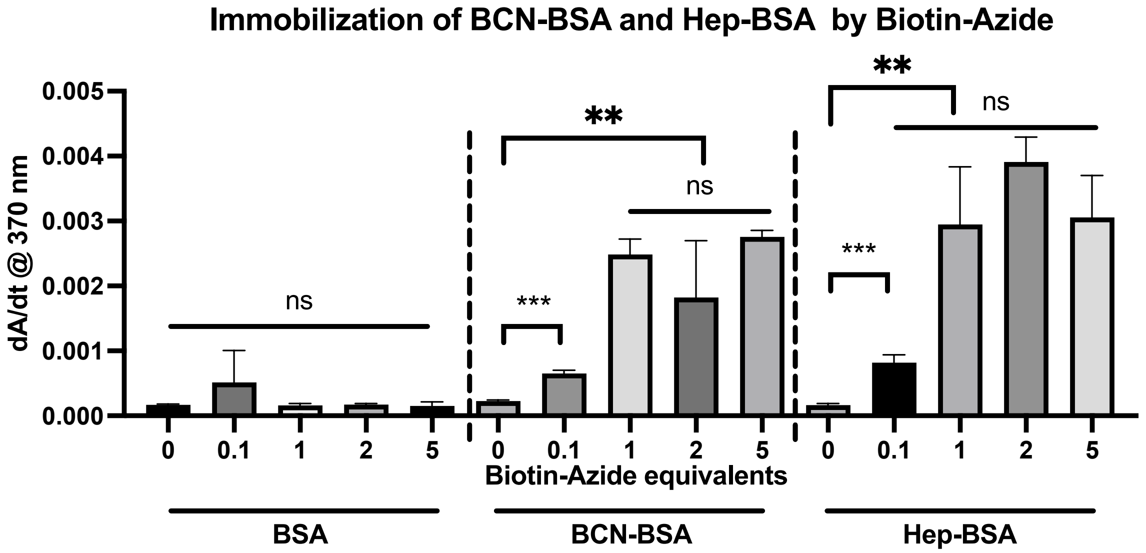

**Figure S12.** Concentration Dependent FGF1 binding to rHS-BSA conjugates

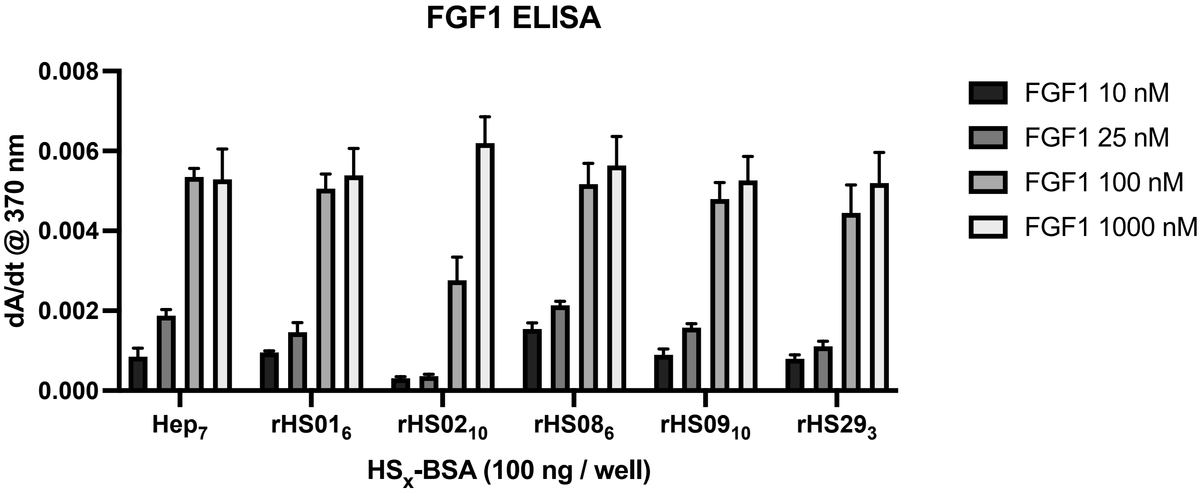

**Figure S13.** Concentration Dependent FGF2 binding to rHS-BSA conjugates

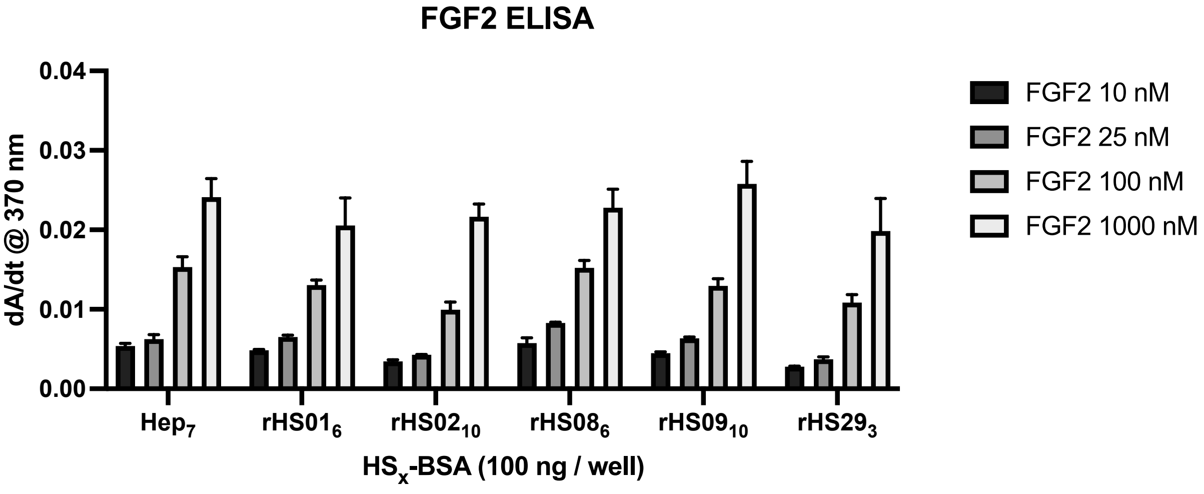

**Figure S14.** FGF2 stimulation of Ext $1^{-/-}$ mESCs in presence of Hep and **Hep$_7$-BSA**

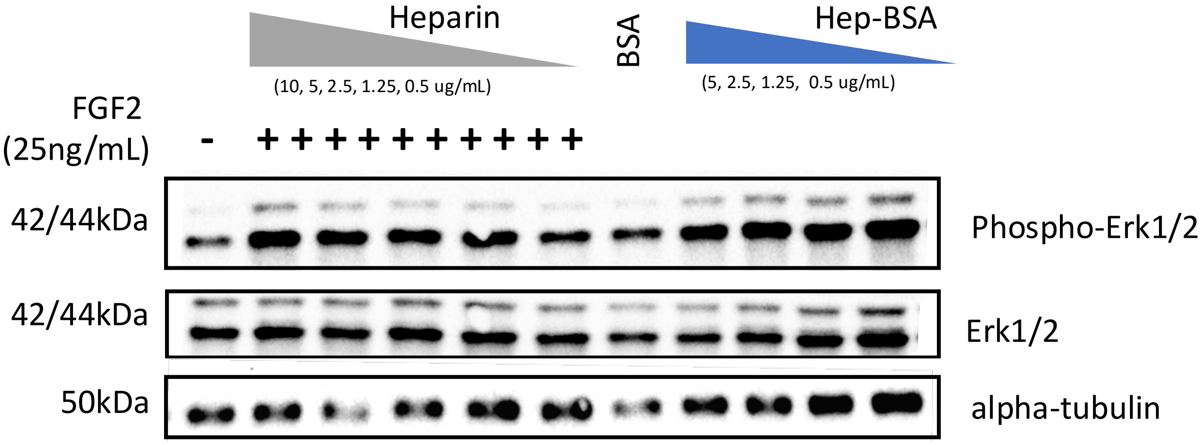

**Figure S15.** FGF2 stimulation of Ext $1^{-/-}$ mESCs in presence of **Hep$_X$-BSA** (x = 1, 2, 4)

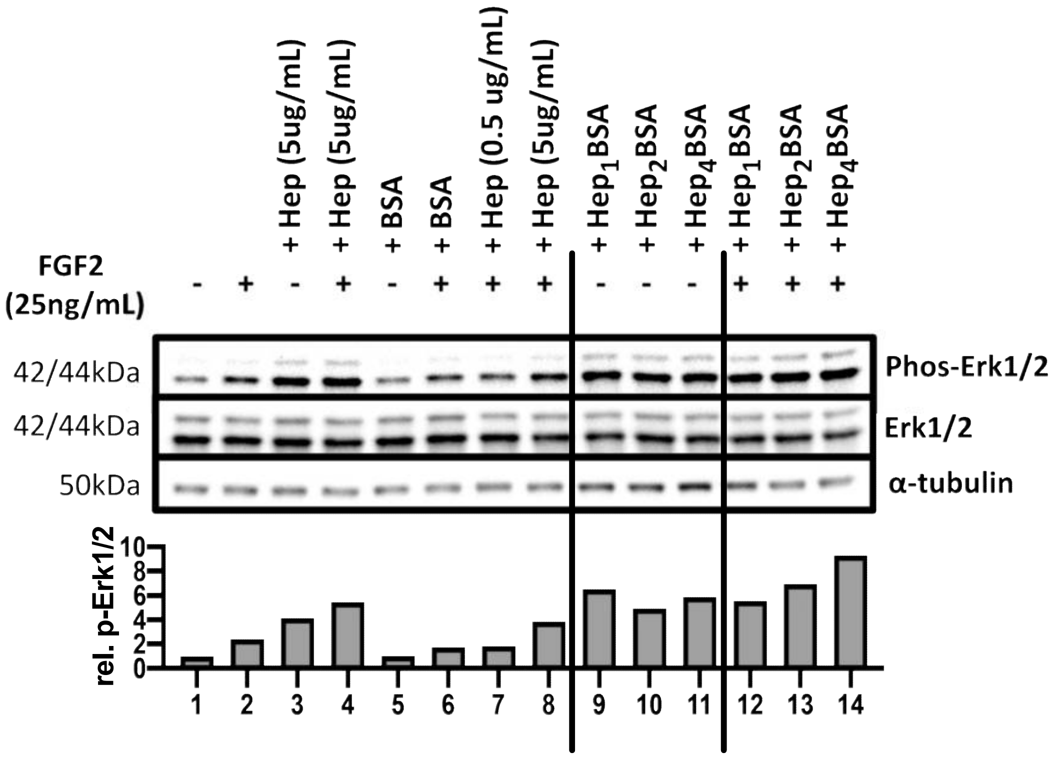
